# Loose ends in cancer genome structure

**DOI:** 10.1101/2021.05.26.445837

**Authors:** Julie M. Behr, Xiaotong Yao, Kevin Hadi, Huasong Tian, Aditya Deshpande, Joel Rosiene, Titia de Lange, Marcin Imieliński

## Abstract

Recent pan-cancer studies have delineated patterns of structural genomic variation across thousands of tumor whole genome sequences. It is not known to what extent the shortcomings of short read (≤ 150 bp) whole genome sequencing (WGS) used for structural variant analysis has limited our understanding of cancer genome structure. To formally address this, we introduce the concept of “loose ends” - copy number alterations that cannot be mapped to a rearrangement by WGS but can be indirectly detected through the analysis of junction-balanced genome graphs. Analyzing 2,319 pan-cancer WGS cases across 31 tumor types, we found loose ends were enriched in reference repeats and fusions of the mappable genome to repetitive or foreign sequences. Among these we found genomic footprints of neotelomeres, which were surprisingly enriched in cancers with low telomerase expression and alternate lengthening of telomeres phenotype. Our results also provide a rigorous upper bound on the role of non-allelic homologous recombination (NAHR) in large-scale cancer structural variation, while nominating *INO80*, *FANCA*, and *ARID1A* as positive modulators of somatic NAHR. Taken together, we estimate that short read WGS maps >97% of all large-scale (>10 kbp) cancer structural variation; the rest represent loose ends that require long molecule profiling to unambiguously resolve. Our results have broad relevance for future research and clinical applications of short read WGS and delineate precise directions where long molecule studies might provide transformative insight into cancer genome structure.

## Introduction

Cancer genomes frequently harbor hundreds of somatic DNA rearrangement junctions, which are pairs of oriented genomic locations that are distant in the reference but adjacent in the tumor genome. Rearrangement junctions contribute to simple (1-2 junctions) or complex (>3 junctions) structural variants (SVs), the vast majority of which also give rise to copy number (CN) alterations (CNAs). Junctions are detected in paired end short read (< 150 bp) whole genome sequencing (WGS) through discordant read mapping,^1–15^ split read alignment,^16–30^ and local assembly.^31–37^ Recent pan-cancer WGS studies have applied novel analytic methods to thousands of short read WGS cancer samples to characterize new classes of complex SVs, including templated insertion events, rigma, pyrgo, and tyfonas.^38, 39^ A broader biological question is how cancer cells tolerate, evolve, and possibly benefit from larger scale (>10 kbp) changes in chromosomal structure.^40–49^

A common perception of short read (< 150 bp) whole genome sequencing (WGS) is that it has limited sensitivity for SV detection, particularly for junctions that arise in repetitive reference sequence.^1, 2, 50–54^ More than two-thirds of the human genome comprises repetitive sequences,^55^ including transposable elements (TEs), satellites (e.g. centromeres), microsatellites, and telomeres; hence, under a model of uniform breakage there is a high probability that a DNA break will occur inside a repetitive element. In addition, specific classes of TEs (e.g. LINEs) have been shown to be active in a subset of cancer types and are associated with somatic insertions^56^ and occasionally rearrangements.^57^ Finally, though cancer DNA repair is thought to mainly involve non-homologous end joining, non-allelic homologous recombination (NAHR) and single-stranded annealing (SSA) may also play a role.^58–62^ The paucity of NAHR and SSA junctions in WGS may be due to technical factors since short read WGS may be unable to map junctions between pairs of similar or identical sequences. While the role of NAHR in constitutional structural variation has been well established, its role in cancer genomes has not been rigorously ruled out.

In the past few years, several studies employing long molecule Pacific Biosciences single molecular real-time (SMRT), Oxford Nanopore Technologies (ONT) nanopore WGS, BioNano optical mapping (OM), and/or 10X Chromium linked read (LR) whole genome profiling have found many examples of SVs that are missed by short-read WGS.^52, 53, 63–73^ However, most of the cancer-associated variants missed by WGS are short-range (< 10 kbp) SVs that resemble large or complex insertions or deletions (indels) rather than large scale structural alterations. These long molecule WGS studies have also been limited to a handful of cell lines, organoids, and/or tumor samples and thus have lacked power to systematically assess global or recurrent patterns of missed variants across many cancer types. As a result, it is still unclear how limits in the SV sensitivity of short read WGS impact our current understanding of cancer chromosomal structure.

A key prerequisite to reasoning about incomplete data is rigorously defining what is missing.^74^ Structural variation has a fundamental coherence which arises from the fact that CNAs and rearrangements form two facets of a single genome structure. This coherence is captured in a data structure called a *junction-balanced genome graph*^39, 50, 71, 75–80^ (genome graph for brevity) which synchronizes the dosage of genomic intervals (i.e. vertices) and their reference or variant adjacencies (i.e. edges) through the application of a junction balance constraint. This constraint applies the notion that every copy of every interstitial genomic segment has a left and a right neighbor, effectively coupling the copy number assignment across rearrangement-induced adjacencies.

We have recently shown that our genome graph inference framework JaBbA^39^ outperforms all previous genome graph and classical (i.e. non graph-based) CNA callers in its ability to characterize genome-wide somatic CNAs. A key aspect of JaBbA’s CNA accuracy is its incorporation of “loose ends”, which represent sites of CNAs that are unaccompanied by a rearrangement junction. While our previous work exploited the accuracy of these graphs to reveal complex SV events, the distribution of loose ends across cancer has not yet been explored. In this study, we analyze JaBbA loose ends to characterize repeat-driven SV events and assess the completeness of large-scale SV reconstructions from short read WGS.

## Results

### Identifying sites of copy number change unexplained by rearrangements

JaBbA fits a probabilistic model to assign integer copy numbers *κ*: *V* ⋃ *E* → ℕ to the vertices *V* and edges *E* of a genome graph *G* = (*V, E*), built from a set of candidate segments and junctions that are provided as input. The *maximum a posteriori* estimate of *κ* maximizes the likelihood of binned-read depth while minimizing the number of loose end edges *L* ⊂ *E* that are given a nonzero copy number subject to a junction-balance constraint (see^39^ for full formulation). This constraint requires the dosage *κ*(*v*) of every vertex *v ∈ V* to be equal to the sum of dosage *κ*(*e*) for all incoming edges *e* ∈ *E*^−^(*v*) (or similarly, outgoing edges *e* ∈ *E*^−^(*v*)). Given this constraint, the model should fit loose ends at all sites of copy number change that are not associated with a rearrangement (**Fig. 1a-b**).

**Fig. 1.**
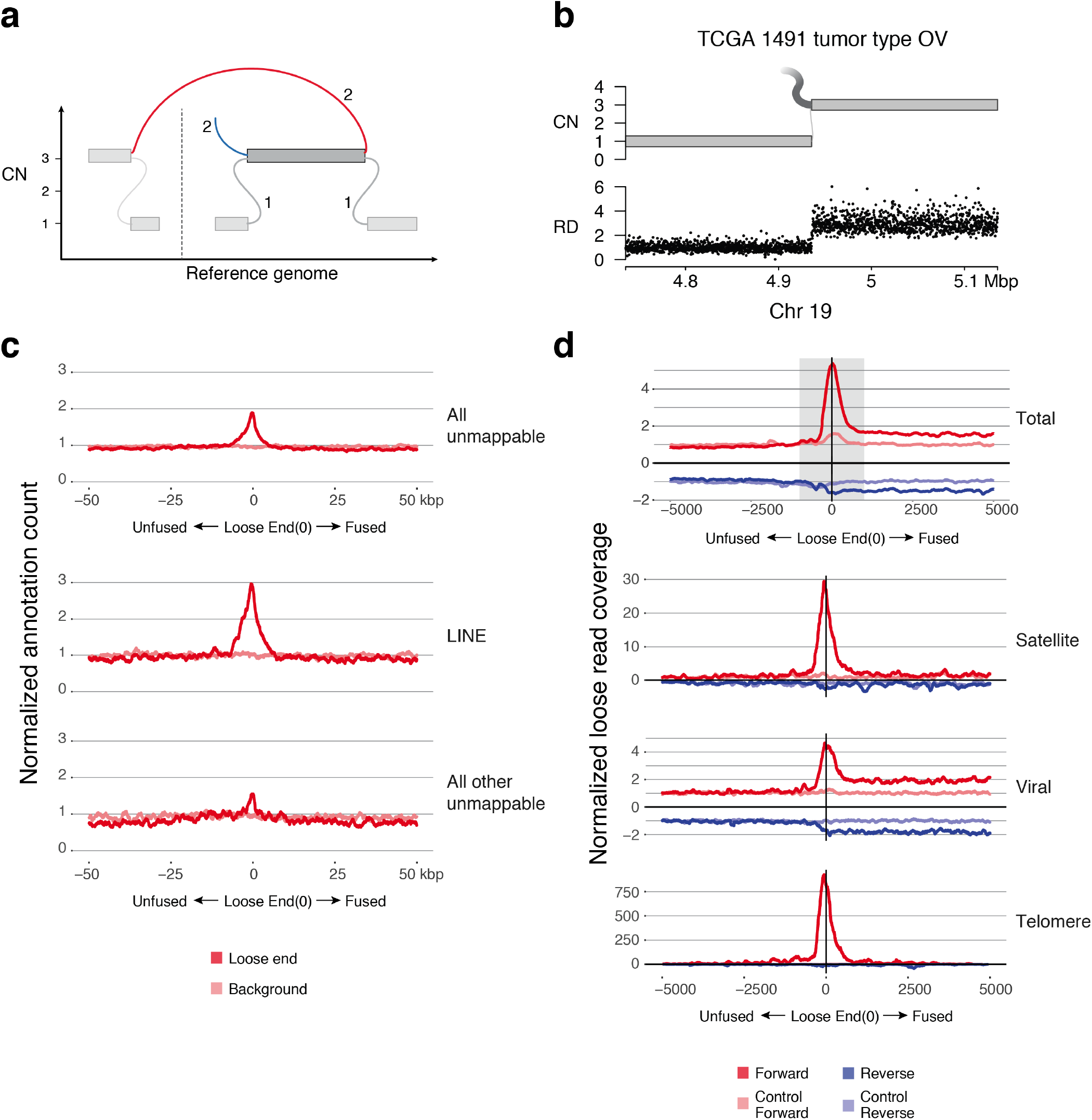
Loose ends are enriched in reference and mated repeats. (a) JaBbA junction balance schematic. The sum of copy numbers of edges + loose ends entering and leaving a node must equal the copy number of the node. Junction shown in red, loose end in blue, reference edge in gray. (b) Example loose end in the graphical representation of TCGA ovarian carcinoma sample 1491. Top, JaBbA graph with a 2-copy loose end. Bottom, normalized binned read depth. (c) Aggregate count of MAPQ=0 annotations surrounding loose ends, normalized to background. Top, aggregate count of all MAPQ=0 coordinates. Middle, aggregate count of MAPQ=0 coordinates overlapping a LINE annotation. Bottom, aggregate count of all non-LINE MAPQ=0 coordinates. Red, aggregate counts around loose ends. Pink, aggregate counts around background sampled coordinates. (d) Aggregate count of loose read alignments surrounding loose ends, normalized to control. Y-axis above zero indicates alignment to loose end-supporting strand; below zero indicates alignment to opposite strand. Top, total aggregate count of all loose reads. Gray box, 1 kbp window around loose end. This window is used throughout analyses to extract aligned reads. Second from top, aggregate count of loose reads whose mates align to satellite annotations. Third from top, aggregate count of loose reads whose mates align to viral sequences. Bottom, aggregate count of loose reads whose mates contain exact matches to 18mers of telomeric motifs. Red, aggregate count from loose end-supporting (“forward”) strand in tumor sample. Pink, aggregate count from forward strand in normal sample. Dark blue, aggregate count from opposite (“reverse”) strand in tumor sample. Light blue, aggregate count from reverse strand in normal sample.

To test this assertion, we generated genome graphs for 2,319 WGS cases spanning 31 tumor types using 200 bp binned tumor and matched normal read depth, CBS segmentation,^81^ and SvABA junction calling^31^ (see Methods). The resulting genome graphs yielded a total of 192,298 large (>10 kbp) junctions (median 50 per sample) and 524,072 candidate loose ends (median 72 per sample). Inspecting read depth patterns across a random sample of these candidate loose ends revealed that a subset occurred at sites of recurrent read-depth “waviness”, which has been previously attributed to fluctuations in GC content^82^ and replication timing-based WGS batch effects^83^ (**Extended Data Fig. 1a-b**). A smaller fraction of candidate loose ends arose from constitutional copy number changes that were also found in the matched normal but were exposed through a somatic aneuploidy that contained the constitutional variant (**Extended Data Fig. 1c-d**).

Following manual review of tumor and matched normal read depth profiles around candidate loose ends, we developed a classifier (see Methods) to exclude false positives. Applying this classifier, we found 25,271 loose ends harboring a *bona fide* copy number change (**Extended Data Fig. 1e**). We manually verified random samples through blinded visual inspection of read depth profiles by five co-authors to establish false negative (19%, 95% credible interval [16.3%, 22.7%]) and false positive rates (2.6%, 95% credible interval [1.8%, 3.6%]) (**Extended Data Fig. 1f**).

### Loose ends are enriched in repetitive and foreign sequences

We postulated that loose ends could arise from unmappable and/or repetitive elements at one or both junction breakpoints. After annotating base level mapping quality scores (MAPQ) through BWA-MEM alignment of short (150 bp) sequence queries generated by applying a sliding window to the hg19 reference (see Methods), we defined the mappable short read genome as any base giving rise to a MAPQ=60 self-alignment. Applying this criterion, we found that only a minority (7.5%) of RepeatMasker annotated features were not mappable (harbored >5% MAPQ<60 bases) (**Extended Data Fig. 2a**). Extending this approach to the full genome, we found that an additional 2.6% and 6.7% of the human reference were unmappable centromere or unannotated unmappable sequence, respectively (**Extended Data Fig. 2b**). These fractions were comparable for 101 bp and 150 bp analyses (**Extended Data Fig. 2b-c**).

**Fig. 2.**
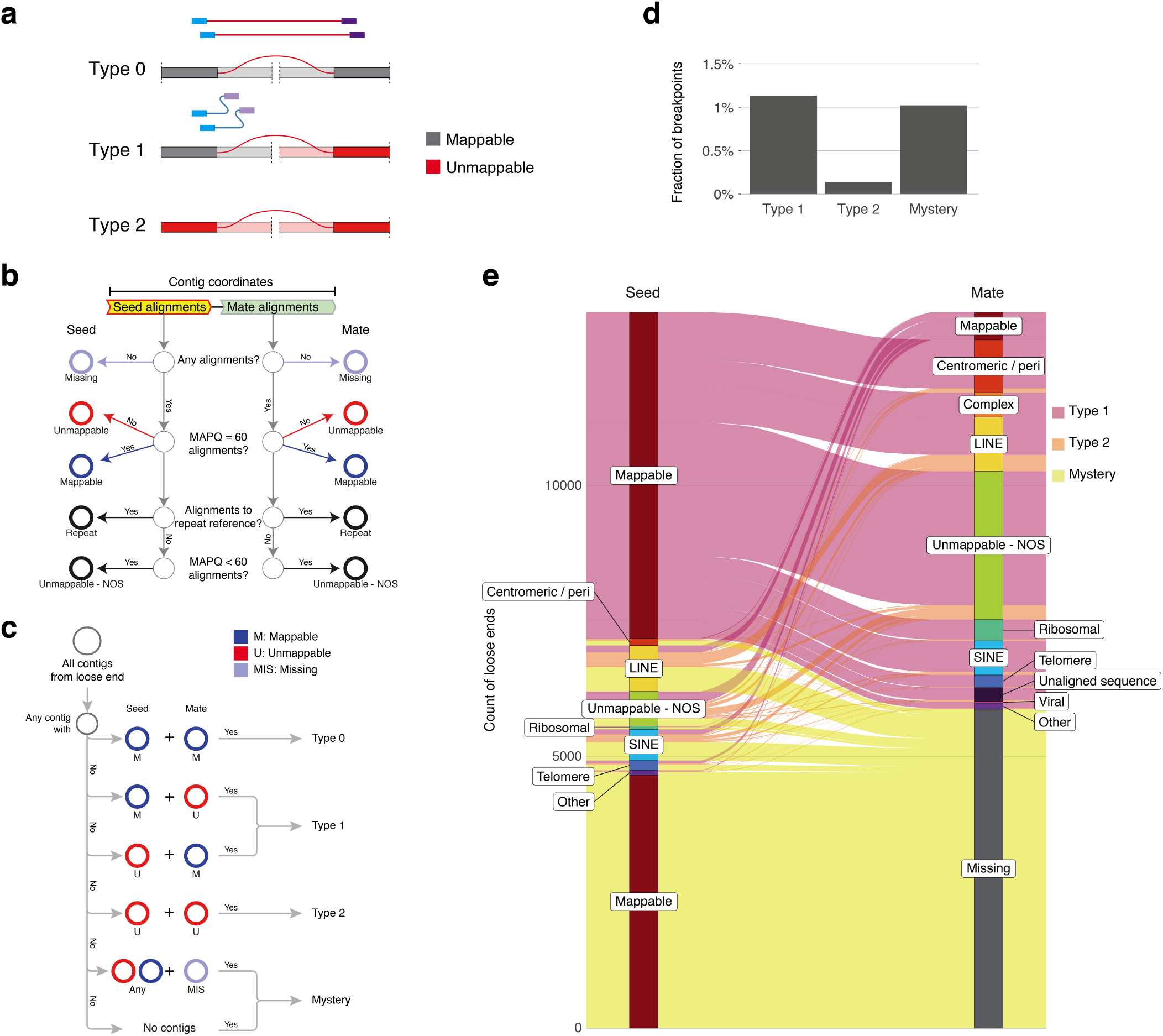
A taxonomy of loose ends. (a) Diagram of mappability contexts for Type 0, 1, and 2 junctions. (b) Flow chart for determining mappability of seed and mate sides of given contig. Every contig side is categorized as mappable or unmappable and separately annotated with the type of foreign sequence, if any, is predominantly present. (c) Flow chart for determining categorization of given loose end, given all assembled contigs. (d) Bar plot of true loose ends falling into each of three categories as a fraction of total SVs in cohort. (e) Alluvial plot of unmappable repeats identified at loose ends.

This analysis revealed that 20.5% of loose ends were within 1 kbp of an unmappable repetitive element, a significant enrichment (*P <* 2.2 × 10^−16^, OR = 1.82) relative to background genomic regions (**Fig. 1c**). This enrichment was particularly strong for LINE elements (*P <* 2.2 × 10^−16^, OR = 2.52). We also found modest enrichment (*P* = 3.96 × 10^−12^, OR = 1.64) of loose ends around all other unmappable regions, including other repeat classes and regions that were not linked to any annotated repetitive elements. These results indicate loose ends preferentially arise in several classes of regions unmappable by short reads. Notably, our analyses did not demonstrate any reference repeat enrichment among candidate loose ends that were removed by our classifier (**Extended Data Fig. 2d**), suggesting that our filtered loose ends encompass the majority of true loose ends.

We next assessed the distribution of sequences mated to reads aligning to loose ends. To do so, we defined a canonical orientation around each loose end by defining a “forward” strand that points in the direction of copy number *decrease* (e.g. left in **Fig. 1b**) and would be expected to harbor alignments with junction-supporting mates. We postulated that high mapping quality (MAPQ=60) reads on the forward strand of loose ends would be enriched in poor-mapping quality (MAPQ=0) mates, which we called *loose reads*. Indeed, we found a distinct tumor-specific enrichment of loose reads in forward but not reverse orientation around mappable loose ends (**Fig. 1d**). Subcategorizing loose reads across those aligning to major classes of repeats, we found profound (>29-fold) enrichment of satellite repeat-associated loose reads within 1 kbp of the loose end relative to background (average loose read count within 5 kbp in control samples), with lower magnitude enrichment (2.9-6.0) observed among loose reads aligning to SINE, LINE, LTR, and simple repeats. (**Extended Data Fig. 2e**).

We also assessed loose reads for the presence of foreign sequences, including viral and telomere repeats. (While telomere repeats are not foreign to human genomes, they are not included in reference genome builds). Indeed, we found that (TTAGGG)n tracts (for n ≥ 3) were strikingly enriched (>900-fold) among forward strand loose reads, but virtually absent on the reverse strand or in control samples (**Fig. 1d**). Analyzing 6251 known viral sequences from the RefSeq viral sequence database, we also found a lower magnitude (4.7fold) enrichment of viral sequences among loose reads on the forward strand (**Fig. 1d**). Taken together, these results indicate that loose ends arise frequently from the fusion of repetitive or foreign sequences.

### A taxonomy of loose ends

To better understand the nature of SVs missed by short read WGS, we developed a taxonomy based on features of sequence alignments and reference locations associated with loose ends. Integrating the above data, we defined three fundamental configurations that rearrangements found in short-read WGS might take: a junction where neither (Type 0), one (Type 1), or both (Type 2) breakpoints are unmappable by short read WGS, (**Fig. 2a**).

Type 0 junctions may give rise to loose ends because of imperfect sensitivity of junction callers, which have been tuned to balance genome-wide precision and recall.^33, 84, 85^ To maximize sensitivity for Type 0 junctions around loose ends, we applied both discordant pair and local assembly analyses (see Methods). Since SvABA already uses local assembly (via String Graph Assembly^86^) we used a de Bruijn graph assembler (Fermi^87^) as an alternative approach to build contigs from junction-supporting reads and their mates (**Extended Data Fig. 3a**). Specifically, we oriented each loose end-derived contig to the forward strand of the loose end and examined features of the seed sequence (the 5’ part of the contig aligning to the loose end locus) and the mate (the distant or novel sequence at the 3’ end of the contig) (**Fig. 2b**). We used features of these chimeric (seed and mate) and distant (mate only) contigs to label each loose end as Type 0, 1, or 2 (**Fig. 2c, Extended Data Fig. 3b**). Loose ends that did not yield a tumor-specific contig with a distant mate alignment were labeled “mysteries” (**Extended Data Fig. 2c, Fig. 3b**).

**Fig. 3.**
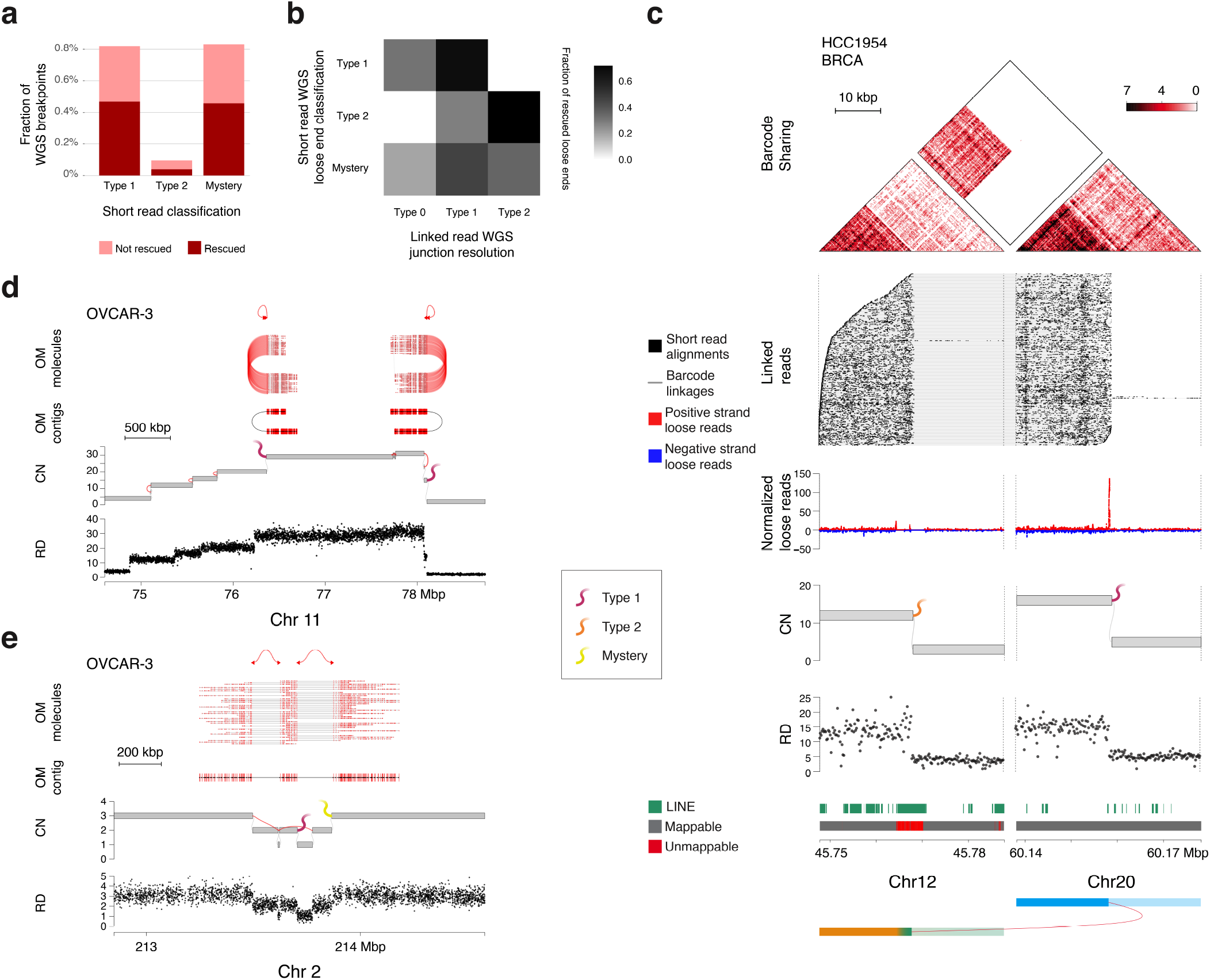
Loose end resolution with linked reads and optical maps. (a) Bar plot of loose ends from long molecule genome profiling cohort categorized by short read WGS analysis. Pink, total loose ends in each category. Red, loose ends rescued by long molecule SV caller. (b) Heatmap of “rescued” categories based on LR sequencing. (c) Example from breast cancer cell line HCC1954 of a pair of loose ends rescued by LR WGS. From top: heatmap showing barcode sharing between 200 bp bins. LR alignments (every horizontal row represents reads sharing one barcode linked by gray lines, with individual read alignments in black). Tumor/normal relative aggregate count of loose read alignments, showing peak at loose end on right. JaBbA graph representation of the sample showing two high copy loose ends corresponding to the apparent junction breakpoints. Normalized binned read depth data, showing corresponding coverage depth change. RepeatMasker LINE annotations, showing 4 kbp LINE overlapping loose end on left. Mappability from simulated 150 bp reads, showing loss of mappability at 4 kbp LINE annotation on left. Diagram of proposed underlying allele. (d-e) Examples from ovarian cancer cell line OVCAR-3 of loose ends rescued by Bionano Genomics OM. From top, rescued junction(s) called from OM Rare Variant pipeline. Alignments of individual OM molecules. Alignment of contig assembled from OM molecules. JaBbA graph with loose ends corresponding to breakpoints of rescued junctions. Normalized binned read depth data, showing corresponding coverage depth change. (d) Two fold back junctions identified by OM within a BFBC pattern. (e) Pair of loose ends joined by OM junction. Left loose end was categorized as Type 1 by short read sequencing. Right loose end was categorized as Mystery by short read sequencing. CN, copy number. RD, read depth.

Applying this approach, we found that almost half (48%, 12,068 of 25,271) of loose ends arose from Type 0 junctions that were missed during genome-wide analysis. These junctions were supported by a contig with a MAPQ=60 chimeric reference genome alignment and/or discordant read-pair cluster supported exclusively by tumor-derived sequences (**Extended Data Fig. 3b**). We propose that these Type 0 breakpoints, combined with those found in the original genome-wide callset, delineate the limit of short read structural variant mappability in these WGS data. The remaining unmappable loose ends, which comprise 2% of all breakpoints in these samples, represent junctions that require long molecule profiling to unambiguously map (**Extended Data Fig. 3c**).

Of these 13,203 unmappable loose ends, 49%, 6%, and 45% were Type 1, 2, and mystery loose ends, respectively (**Fig. 2d**). To categorize unmappable loose ends further, we labeled seed and mate regions of loose ends with fifteen classes of repeats and/or foreign (e.g. viral) sequences (**Fig. 2e**, **Extended Data Fig. 3d**, **Extended Data Fig. 4**). Type 1 loose ends were most frequently the result of LINE, SINE, telomeric, and peri-centromeric sequences that were connected to a mappable seed region. A subset (792, 12%) of Type 1 loose ends yielded contigs with multiple mappable (MAPQ=60) but complex multi-part alignments whose precise order and orientation could not be unambiguously resolved, indicating that these were the result of complex rearrangements that short read WGS could only partially reconstruct (**Extended Data Fig. 3d**). Type 2 loose ends were most frequently the result of fusions of reference non-adjacent LINE elements and pairs of unannotated low mapping quality regions. While mysteries most frequently comprised mappable seed regions, they were also frequently associated with unmappable repetitive elements, including LINE, SINE, peri-centromeric, and telomeric sequence, suggesting that these loci may harbor additional Type 2 loose ends.

**Fig. 4.**
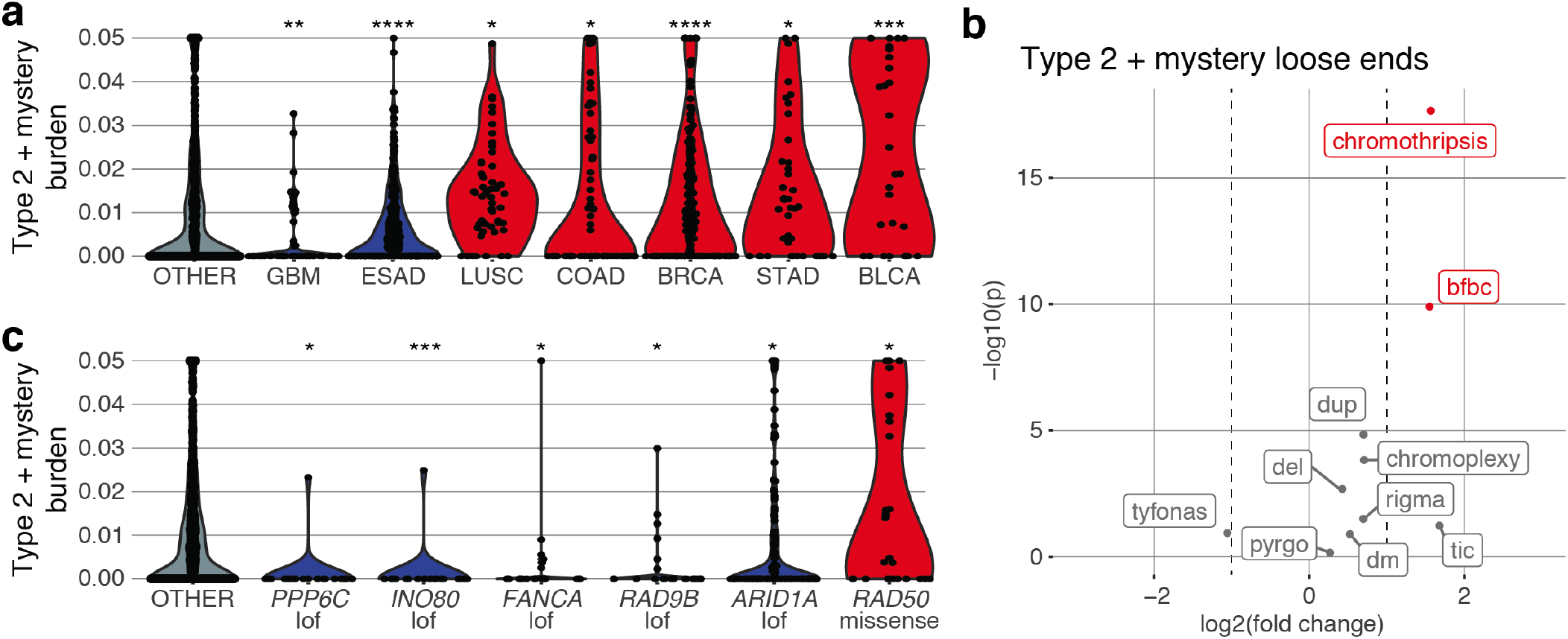
Bounding the burden of NAHR in cancer genomes. (a) Violin plots of tumor types significantly associated with Type 2 + mystery loose ends, which bound the burden of non-allelic homologous recombination in cancer genomes. Blue, tumor type significantly depleted in Type 2 + mystery loose ends. Red, tumor type significantly enriched in Type 2 + mystery loose ends. Y-axis, fraction of breakpoints per sample categorized as Type 2 or mystery. (b) Volcano plot of association between complex SV patterns and Type 2 + mystery loose ends. (c) Violin plots of genotypes significantly associated with Type 2 + mystery loose ends. Each dot is one tumor. Blue, genotype significantly depleted in Type 2 + mystery loose ends. Red, genotype significantly enriched in Type 2 + mystery loose ends. Y-axis, fraction of breakpoints per sample categorized as Type 2 or mystery. Significance levels: **** fdr < 1 × 10^*−*4^, *** 1 × 10^*−*4^ < fdr < 1 × 10^*−*3^, ** 1 × 10^*−*3^ < fdr < 0.01, * 0.01 < fdr < 0.1.

### Long molecule loose end rescue

Long molecule genome profiling technologies including LR WGS and OM employ the alignment and/or assembly of long (>20 kbp) molecules to map structural variants, including those arising in repetitive genomic regions.^88, 89^ We reasoned that these approaches would be particularly useful for resolving of Type 1 and 2 junctions around unmappable loose ends. To assess this, we generated LR WGS (median molecule size ~50-80 kbp, genome-wide physical coverage 170-215x) and Bionano OM data (median molecule size ~310-375 kbp, genome-wide physical coverage 400-430x) for 10 and 3 cancer cell lines, respectively (see Methods).

We examined the density of SV calls from three standard LR algorithms (GROC-SV,^65^ LinkedSV,^90^ and NAIBR^91^) and the Bionano Solve Rare variant pipeline (see Methods) in the vicinity of loose ends. We found that loose ends were markedly enriched in both LR and OM-derived junction breakpoints at a distinct peak within 1 kbp (LR) or 10 kbp (OM) oriented to the loose end forward strand (**Extended Data Fig. 5a**). Long molecule analysis resolved junctions for 54%, 41%, and 56% of Type 1, 2, and mystery loose ends (**Fig. 3a**). Of note, we found a somewhat lower rate of validation using LR (46%, 44%, and 49% of Type 1, 2, and mystery loose ends, respectively) than OM (73%, 33%, and 70%).

**Fig. 5.**
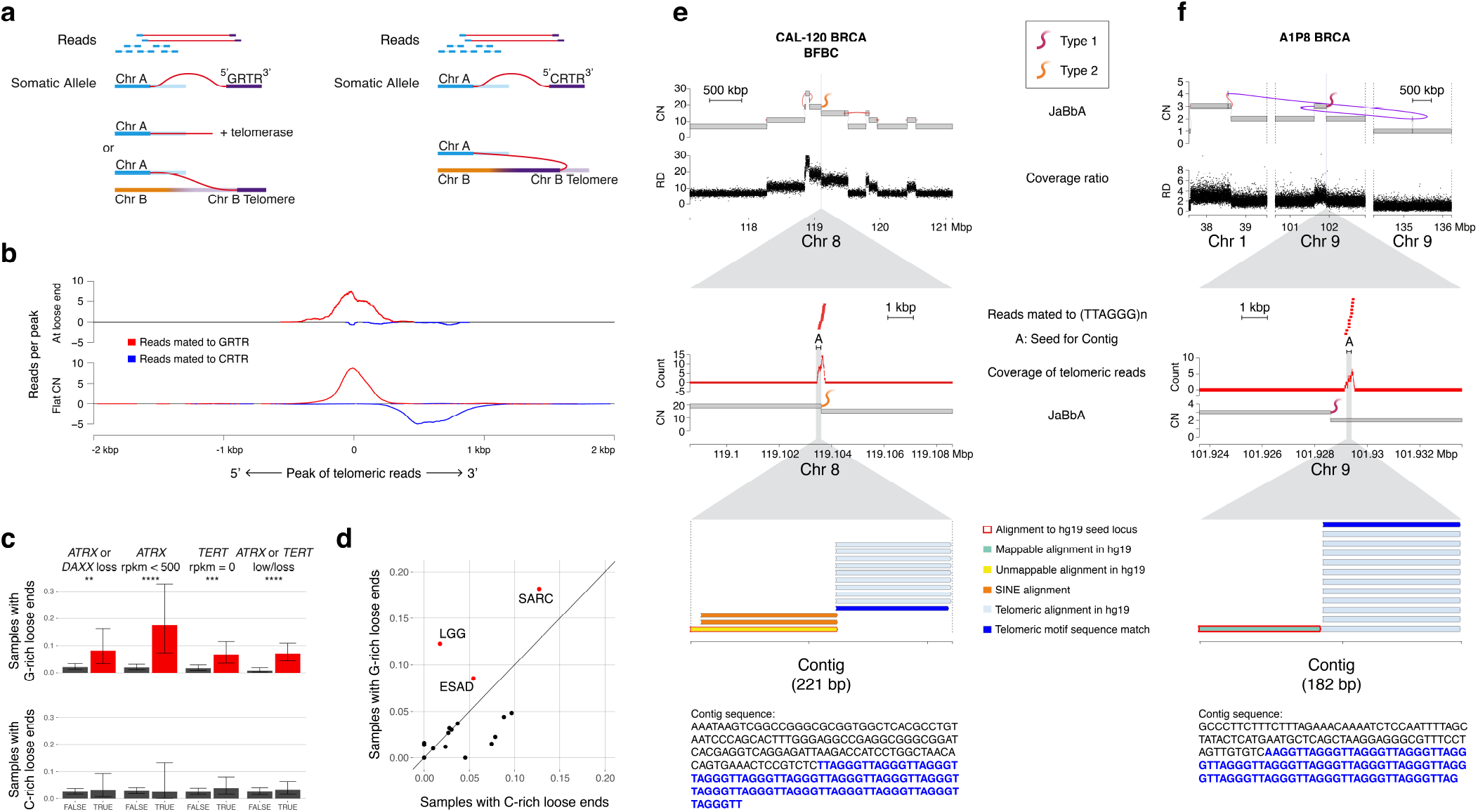
G-rich telomere repeats nominate new cancer chromosome ends. (a) Diagram of proposed telomere-involved rearrangements. At a chromosome terminus, the loose end strand will be mated to telomeric (TTAGGG)n repeats. At a rearrangement within a telomere oriented away from the chromosome terminus, the loose end strand will be mated to (CCCTAA)n repeats. (b) Aggregate alignments of telomeric mates with respect to the strand and position of a G-rich telomeric mate peak, normalized to total count of peaks contributing to each track. Top, telomeric mate peaks ≤ 1 kbp away from a loose end. Bottom, telomeric mate peaks > 1 kbp away from a CNA. Red, reads mated to G-rich telomeric repeats. Blue, reads mated to C-rich telomeric repeats. Positive y-axis, reads aligned to same strand as G-rich telomeric mate peak. Negative y-axis, reads aligned to opposite strand as G-rich telomeric mate peak. (c) Bar plot of fraction of samples with a given genotype containing at least one telomeric loose end. Error bars show 95% confidence interval. Red, significant enrichment of telomeric loose ends within genotype. Gray, not significant. “TRUE”, samples with the genotype. “FALSE”, samples without the genotype. Significantly enriched genotypes (compared to samples without the genotype) are marked by asterisks (Fisher’s exact test). Significance levels: **** (*P <* 1 × 10^*−*4^), *** (*P <* 1 × 10^*−*3^), ** (*P <* 0.01), * (*P <* 0.05). (d) Scatter plot of fraction of samples containing at least one C-rich telomeric loose end vs fraction of samples containing at least one G-rich telomeric loose end separated by tumor type. Red, tumor type significantly enriched for G-rich telomeric loose ends. Gray, not significant. (e) Example of a neo-telomere from breast cancer carcinoma cell line CAL-120 which falls within a BFBC annotation. Top, 4 Mbp window around neo-telomeric loose end, showing nearby junctions and coverage changes. Middle, 10 kbp window around neo-telomeric loose end, showing individual alignments of reads mated to telomeric reads, location of the seed region used as input to assembly pipeline, and aggregate count of strand-specific telomeric read alignments. No telomeric reads are found in this window on the reverse strand or in the paired normal sample. Bottom, representative contig output from assembly pipeline. The seed is unmappable with corresponding SINE alignments, and the mated sequence is exact matches to (TTAGGG)n. Representative contig sequence shown below. Black, sequence aligning to seed region. Blue, mated sequence. (f) Example of a neo-telomere from TCGA breast cancer carcinoma sample A1P8. Panels same as (e). Top, neo-telomeric loose end falls within templated insertion chain pattern. Bottom, seed is mappable. CN, copy number. RD, read depth.

We next probed the fate of resolved loose ends as a function of their short read WGS-derived classification. Specifically we classified junctions inferred through long molecule profiling as Type 0, 1, 2 on the basis of the short read mappability of the fused reference region. Since OM profiles provide approximate breakpoint coordinates (~20 kbp resolution), we limited this analysis to LR WGS (**Fig. 3b**). While the majority (68.6%) of LR-resolved Type 1 loose ends yielded Type 1 junctions, we found a subset (31.4%) that were resolved into Type 0. Similarly, while a majority (71.4%) of resolved Type 2 loose ends yielded Type 2 junctions, a subset (28.6%) of these were resolved into Type 1 junctions. Finally, we found that mystery loose ends were resolved into all three junction classes, including a substantial fraction to Type 2 (35.4%).

Integrating these results, we found that the majority (77.3%) of loose ends that were resolved into Type 2 junctions by LR WGS were classified as mystery loose ends in the short read WGS analysis. Furthermore, the overall frequency of Type 2 junctions in LR WGS (0.84%) was similar to the combined rate of Type 2 or mystery loose ends (1.2%) in our short read analyses. As a result, we conclude that the rate of Type 2 and mystery loose ends in short read WGS provides an upper bound on the rate of Type 2 junctions in cancer genomes.

To better understand the re-classification of Type 1 and 2 loose ends by long molecules we examined a pair of differently classified loose ends in the metastatic breast cancer cell line HCC1954 (**Fig. 3c**) that were fused into a Type 1 junction following LR profiling. The Type 1 loose end at chromosome 20 resides in a mappable locus harboring a large peak of loose reads indicating fusion to an unmappable LINE repeat. The loose end on chromosome 12 was located in a poorly mappable LINE element and associated with loose reads mapping to a distant and also unmappable LINE. LR WGS confirmed support for a high copy junction joining one mappable and one unmappable breakpoint, supporting a Type 1 junction classification. Re-alignment of reads from junction-supporting vs. reference LR barcodes indicated the presence of junction-specific sequences mapping to loose end-associated contigs derived from short read WGS. These results are consistent with the fusion of a mappable chromosome 20 locus to an unmappable reference LINE on chromosome 12 with the insertion of a second unmappable LINE element at the breakpoint.

We next inspected the impact of LR and OM loose end resolution on complex SV inference. We found two loose ends associated with a breakage fusion bridge cycle (BFBC) on chromosome 11 in ovarian cancer cell line OVCAR-3 profiled with OM and LR WGS. Both were categorized as Type 1 loose ends with multiple ambiguous mate alignments. The leftmost loose end additionally harbored inserted LINE sequence between the two fused sides. According to our recently published genome-graph complex SV taxonomy,^39^ this event was classified as a BFBC because the total junction copy number across the five fold-back inversion junctions was nearly equal to the height of this relatively simple amplicon. Analyzing these genomes with OM, we found fold-back inversion junctions at both loose ends. While these findings did not change the complex SV classification, they substantially strengthened the fold-back inversion signal supporting the BFBC classification (**Fig. 3d**).

In the same cell line, we found two loose ends at a chromosome 2 locus also harboring a pair of Type 0 deletion junctions on short read WGS (**Fig. 3e**). The Type 1 loose end was associated with mappable seed mated to unmappable sequence after short read assembly. The loose end on the right was characterized as a mystery, with unmappable SINE sequence found at a seed that yielded no chimeric contigs. OM analysis revealed a deletion-like Type 1 deletion junction, on the same allele as one of the Type 0 deletions detected with WGS. Following the addition of this junction and re-analysis of the graph, this cluster of deletions was called a rigma, a class of complex SVs revealed in our recent pan-cancer analysis of genome graphs.^39^

### Pan-cancer estimates of non-allelic homologous recombination in cancer genomes

NAHR is a DNA repair mechanism that joins distant but nearly identical sequences; NAHR-driven structural variants will therefore yield Type 2 junctions and be underdetected in short read WGS. Having established a taxonomy of loose ends, we reasoned that the burden of Type 2 and mystery loose ends may provide a proxy for Type 2 junctions and hence NAHR in cancer genomes. While Type 2 junctions appear rare in cancer genomes (1% of all breakpoints), we observed an order of magnitude variation in their burden (0.31%-2.64%) across cancer, with five tumor types (bladder (BLCA), breast (BRCA), stomach adenocarcinoma (STAD), colon adenocarcinoma (COAD), and lung squamous cell (LUSC)) showing significant enrichment relative to their complement (FDR<0.1) (**Fig. 4a, Extended Data Fig. 5b**). Tumor types enriched for Type 2 and mystery loose ends were distinct from those enriched in Type 1 loose ends, which included head and neck squamous cell (HNSC), malignant lymphoma (MALY), and sarcoma (SARC) (**Extended Data Fig. 5c**). Notably, breast cancer was significantly enriched in Type 2 and mystery but depleted for Type 1 loose ends.

To explore the possible impact of NAHR on SV mutational processes, we examine the overlap of Type 2 and mystery loose ends with distinct classes of complex SVs identified through the analysis of genome graph topology^38, 39, 92^ (**Fig. 4b, Extended Data Fig. 5d**). We found the most significant overlap of Type 2 and mystery loose ends with chromothripsis (*P* = 2.24 × 10^−18^, RR = 2.97, Wald test) and BFBCs (*P* = 1.26 × 10^−10^, RR = 2.93). In contrast, Type 1 loose ends were significantly enriched at double minutes (*P* = 1.10 × 10^−24^, RR = 4.34) and templated insertion chains (*P* = 3.01 × 10^−3^, RR = 4.75). These results indicate differential mutational processes generating Type 1 and 2 junctions in cancer genomes.

We hypothesized that acquired defects in DNA repair may modulate the rate of NAHR in mutant tumors. To investigate this, we correlated the burden of Type 2 and mystery loose ends with the presence of alterations at 191 frequently mutated (>1% of cases) DNA damage response genes.^93^ After correcting for tumor type, this analysis identified six genes (FDR < 0.1) linked with enrichment or depletion of Type 2 and mystery loose ends (**Fig. 4c, Extended Data Fig. 5e**). The most significant association among these was with *INO80*, a nucleosome remodeling enzyme that has been implicated in double stranded break (DSB) motility and mitotic homologous recombination.^94–100^ Tumors harboring somatic truncating mutations in *INO80* showed a significant depletion in the burden of Type 2 and mystery loose ends relative to wild type tumors (*P* = 7.18 × 10^−7^, RR = 0.082). We found a similar depletion (*P* = 2.13 × 10^−3^, RR = 0.647) in tumors with truncating mutations of *ARID1A*, a major component of the SWI/SNF nucleosome remodeling complex.

Additional negative associations (*P* = 1.38 × 10^−3^, RR = 0.324) included *FANCA*, a member of the Fanconi anemia interstrand crosslink repair complex that has been implicated in homology driven strand-annealing following double stranded DNA breakage,^101^ and two less well characterized regulators of the G1-S cell cycle checkpoint (*PPP6C*, *RAD9B*). The only positive association was with *RAD50*, a component of the MRE11-RAD50-NBS1 (MRN) complex that drives the initial processing of DSBs in both NHEJ and homologous recombination (HR). Interestingly, this association was strongest with tumors harboring missense mutations in *RAD50* (*P* = 2.47 × 10^−3^, RR = 2.14), consistent with these mutations shifting the NHEJ / HR balance rather than abrogating MRN complex function. Taken together, these results provide some of the first cancer genomic evidence to nominate specific DNA repair factors as modulators of cancer NAHR.

### G-rich telomere repeat positive loose ends in telomerase low tumors

While loose ends primarily represent the technical frontier of short read SV mapping, some may represent true novel cancer chromosome “ends”. In addition, the high enrichment of telomere repeats among loose reads (**Fig. 1d**) and contigs associated with Type 1 loose ends in our taxonomy (2.1%, **Fig. 2e**) suggested that some loose ends may represent examples of telomere deposition at novel chromosome ends. As illustrated in (**Fig. 5a**), we predict neotelomeres to yield hotspots of G-rich telomere repeat (GRTR) sequences in loose reads and loose end-derived contigs when oriented to the forward loose end strand. In contrast, C-rich telomere repeat (CRTR) loose end contigs are not consistent with neotelomeric loose ends, but more consistent with telomeric insertions or end-to-end fusion (**Fig. 5a**)

To test this intuition, we compared the distribution of mates with GRTR+ and CRTR+ reads within and outside of the context of loose ends (**Fig. 5b**). Telomere insertions should not result in a copy change and should harbor reciprocal peaks connecting opposite strands of the inserted telomere repeat to the flanking reference regions. Consistent with this prediction, we found that hotspots of reads with GRTR+ mates outside of the context of loose ends were usually associated with a reciprocal peak of opposite strand reads harboring CRTR+ mates an insert size (~600-700 bp) downstream. In contrast, the same analysis performed on GRTR+ loose ends did not reveal a reciprocal peak, suggesting that these events were not the result of telomere insertions.

We next hypothesized that high levels of telomerase activation, such as that associated with *TERT* amplification might be associated with a high neotelomere burden, due to telomerase-mediated double strand break healing.^102^ Comparing loose end burdens in various somatic genetic contexts of telomerase activation, we did not find a difference in the GRTR+ burdens between high-level (CN > 2 × ploidy) *TERT* amplification or over-expression (z-score > 2) (5.6%, *P* = 0.058 and 4.5%, *P* = 0.336 respectively) relative to samples with wild type and average *TERT* expression (**Extended Data Fig. 5f**).

Contrary to our hypothesis, we found a striking enrichment of GRTR+ loose ends in samples with low or negligible *TERT* expression (z-score < −2). There was no difference in the burden of CRTR+ loose ends among any of these *TERT* tranches (**Fig. 5c**)). Since tumors that fail to express telomerase may employ the alternative lengthening of telomeres (ALT) pathway, we asked whether neotelomere burden correlated with loss of function mutations in the key ALT suppressors *ATRX* or *DAXX*.^103^ Indeed we found a higher burden of GRTR+ loose ends in *ATRX* / *DAXX* null tumor samples (**Fig. 5c**). Furthermore, we found that several ALT-associated cancers, including sarcomas (18%, OR = 6.47, *P* = 1.95 × 10^−5^) and low grade gliomas (12.3%, OR = 3.92, *P* = 4.1 × 10^−3^), had the highest rate of GRTR+ loose ends relative to other tumor types **Fig. 5d**).

To yield stable derivative chromosomes, complex SVs like chromothripsis and BFBCs must acquire a telomere or circularize. Telomere acquisition can occur through fusion to an existing chromosome end or through *de novo* telomere synthesis. Investigating the latter possibility, we found instances of GRTR+ loose ends falling within or near the footprints of complex SVs. This included a BFBC in breast cancer cell line CAL-120 harboring a GRTR+ loose end (**Fig. 5e**). GRTR+ loose reads were found on the loose end-supporting strand only, all aligning within 200 bp of the loose end. The assembled contig contained 123 bp of the loose end locus fused to 98 bp of GRTR sequence. This pattern was consistent with a breakage fusion bridge process terminating through neotelomere synthesis. We found another neotelomere in the TCGA breast cancer sample A1P8 amid a cluster of rearrangements between chromosomes 1 and 9 (**Fig. 5f**). This loose end was classified as Type 1, with a mappable reconstruction of the loose end locus again fused directly to 98 bp of GRTR+ sequence. The pattern is consistent with a templated insertion chain^38, 39^ joining chromosome 1 and 9 before terminating in a downstream neotelomere.

### Viral loose ends resolve high-copy amplicons

A subset of cervical (CESC), oropharyngeal squamous (HNSC), and liver hepatocellular carcinoma (LIHC) arise in the context of chronic infection with double stranded DNA viruses (Human Papilloma Virus, HPV; Epstein Barr Virus, EBV).^104–107^ While somatic viral integrations into the host genome are routinely detected by many SV pipelines (e.g. SvABA, GRIDSS2, Aperture),^31, 33, 108^ these algorithms do not readily distinguish between small insertional and large-scale structural variants. Since our genome graphs are defined across the human reference genome, large-scale unbalanced viral SVs would be predicted to result in loose ends (**Fig. 6a**).

**Fig. 6.**
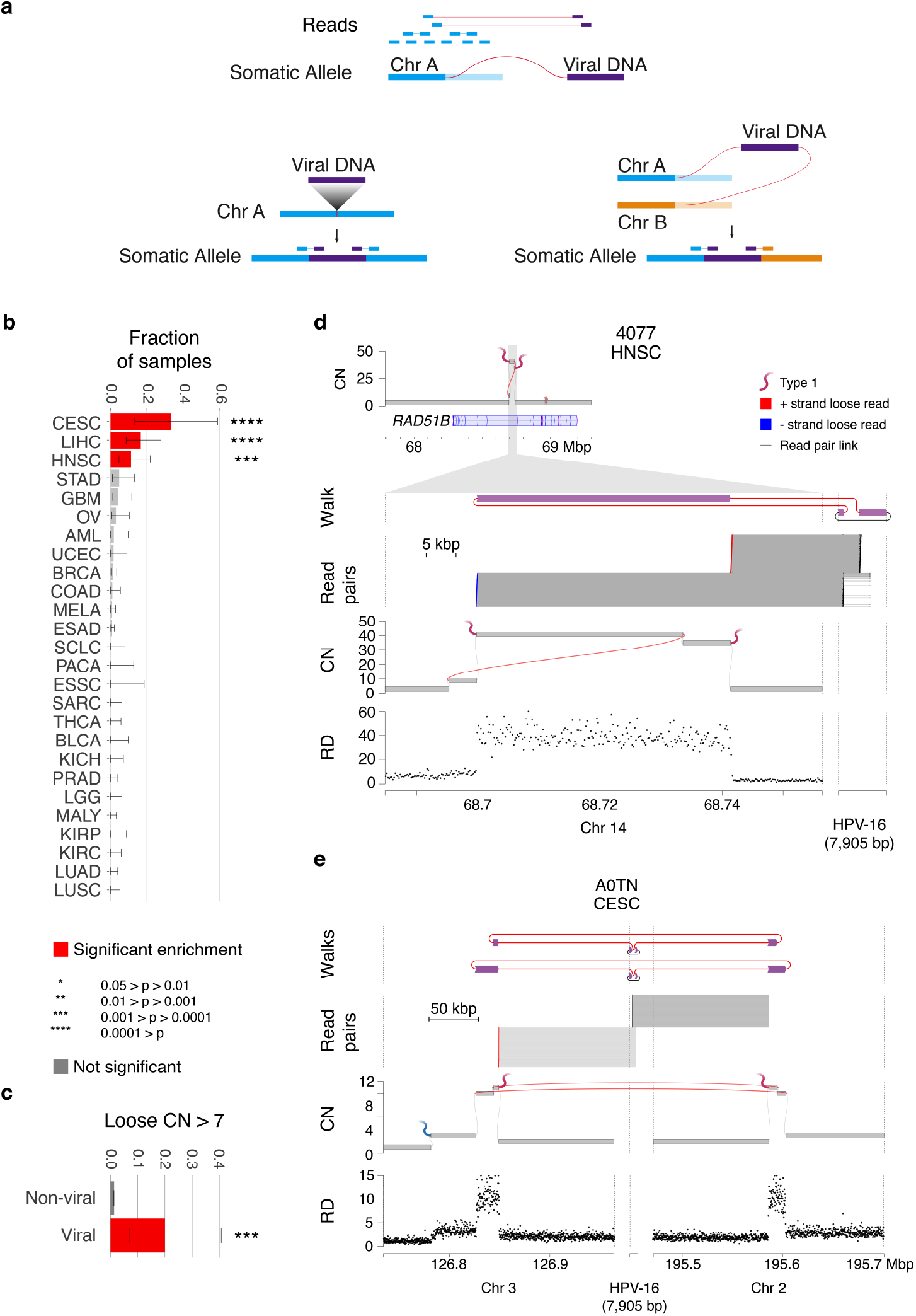
Viral loose ends are amplified in virus-driven cancers. (a) Diagram of expected read sequences given proposed virus-involved rearrangements. A viral insertion will not result in a CNA. A rearrangement mediated by inserted viral sequence will result in a pair of loose ends both mated to viral sequences. (b) Bar plot of fraction of samples containing at least one viral breakpoint (loose end or viral SV call from SvABA) separated by tumor type. Error bars show 95% confidence interval. Red, tumor type significantly enriched for viral breakpoints (Fisher’s exact test). Gray, not significant. (c) Bar plot of fraction of loose ends with a high (> 7) loose copy number. Viral loose ends are significantly enriched in high loose copy number compared to all other loose ends (*P* = 1.88 × 10^*−*5^, OR = 18.6). Error bars show 95% confidence interval. (d-e) Examples of apparent viral-mediated rearrangements. From top, “walk” representation of proposed circular allele. Alignments of read pairs with discordant alignments between human and viral genome. JaBbA graph representation of the sample showing two loose ends corresponding to the discordant read alignments. Normalized binned read depth data, showing corresponding coverage depth changes. CN, copy number. RD, read depth. (d) Example from TCGA head and neck squamous cell carcinoma sample 4077. Top left, zoomed out view of amplicon showing overlap with intron of *RAD51B*. (e) Example from TCGA cervical carcinoma sample A0TN.

Investigating viral contributions to loose ends across our taxonomy (**Fig. 1d**, **Fig. 2e**), we identified 29 Type 1 loose ends that harbored a viral mate, 13 and 9 of which were linked to HPV and HBV, respectively. Though these events were rare (occurring in 0.95% of cases) across cancer, they were significantly more frequent in CESC (*P* = 5.15 × 10^−7^, OR = 29.5), HNSC (*P* = 1.22 × 10^−4^, OR = 7.62), and LIHC (*P* = 1.55 × 10^−8^, OR = 13.4) as well as cases previously annotated as being virus positive (**Fig. 6b**). These results indicate that viral driven SVs are a frequent feature in cancers that arise in the context of viral infection.

Each loose end, like each junction and interval, is associated with a dosage or copy number. Analysis of dosage at 29 viral loose ends showed that these had a significantly elevated copy number relative to non viral loose ends (*P* = 1.7 × 10^−4^, OR = 8.66), **Fig. 6c**). Investigating these high copy viral loose ends, we found a chromosome 14 double minute in an HNSC tumor (TCGA-4077) where two high copy Type 1 loose ends mapped to opposite ends of the HPV-16 genome (**Fig. 6d, Extended Data Fig. 5g**). These loose ends flanked an intronic region of the *RAD51B* gene that was amplified to 40 copies. The most likely reconstruction of this locus was an amplified circular structure with HPV-16 linking the two amplicon ends as an insertion in a duplication-like junction, consistent with a double minute.

We found a similar pattern in a more complex amplicon in CESC case (TCGA-A0TN) (**Fig. 6e**). This subgraph harbored two junctions connecting two distinct foci of increased copy number (CN 10-12) in an intergenic region on chromosome 2. As with the HNSC case, two Type 1 loose ends were linked to opposite sides of the HPV-18 genome, consistent with a high copy junction linking chromosomes 2 and 3 with a viral sequence inserted at the junction (**Extended Data Fig. 5h**). Re-analysis of this genome graph after addition of this inferred high-copy translocation junction enabled reclassification of the associated subgraph as a double minute.

Analysis of additional viral loose ends also revealed examples of simpler and low copy intra- and inter-chromosomal rearrangements. For example, a pair of loose ends on chromosome 8 were linked to opposite ends of the HPV-18 genome in a CESC case (TCGA A2RM) in a tandem duplication orientation (**Extended Data Fig. 6a**). Two additional examples of low copy EBV-linked loose ends were found in a LIHC cell line (SNU-475) and primary tumor sample (TCGA-A5NP) (**Extended Data Fig. 6b-c**). Each pair of loose ends which mapped to opposite ends of the EBV genome could thus be resolved into an inter-chromosomal junction likely representing a simple translocation event.

### Missing structural variants at cancer drivers

A key goal of cancer genome sequencing is to characterize biologically important or actionable driver alterations at known cancer genes. However, it is largely unknown to what degree alterations involving repetitive or foreign sequence contribute to SVs at clinically important cancer drivers. Missing junctions can prevent the detection of a protein-coding DNA fusion or the incorrect grouping of copy number alterations into complex SV events using short read WGS. Across 2,319 cases, we found loose ends contributed to only 5 of a total of 321 breakpoints in the gene bodies of *ABL*, *ALK*, *RET*, *ROS*, *MET*, *BRAF*, *FGFR1/2/3*, and *NTRK1/2/3*, indicating that protein coding fusions at these clinically important loci are rarely missed by short read WGS.

To assess the contribution of loose ends to recurrent copy number alterations at cancer driver genes, we focused on 99 COSMIC Cancer Gene Census genes that have been previously implicated in recurrent peaks of copy number gain or loss in cancer (including 4 genes associated with both) using the GISTIC algorithm.^109^ We then used analyses of local genome graph topology to associate regions of gain (copy number > twice ploidy) or loss (copy number ≤ 1) with specific loose ends (see Methods). Among the 46 recurrently amplified oncogenes, *CCND1* and *MYC* were the most frequently amplified by a loose end (12 and 9 instances respectively), while *BCR* and *MYCL* had the highest fraction of amplifications associated with a loose end (11.5% and 11.4% of total amplifications respectively) (**Fig. 7a**).

**Fig. 7.**
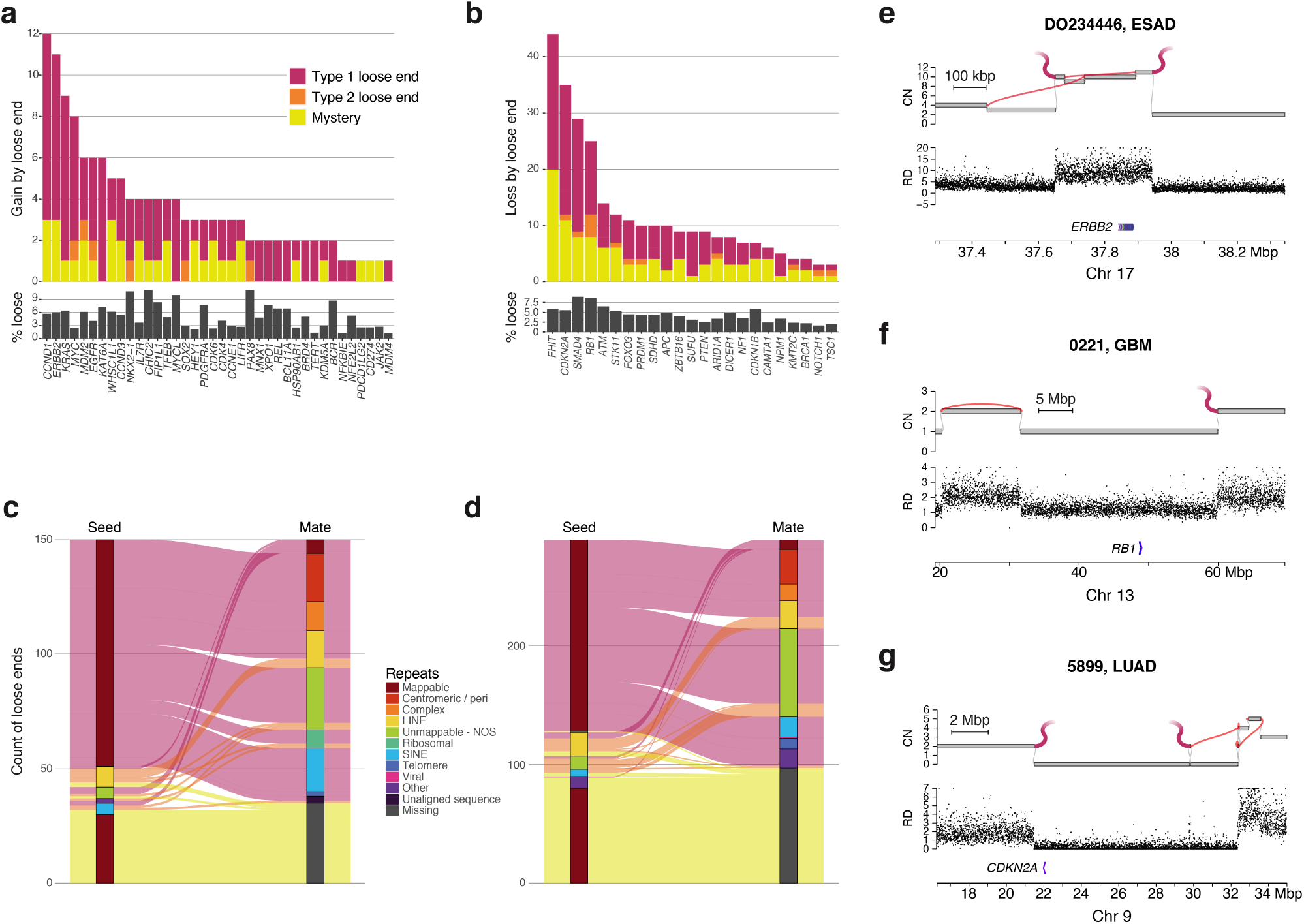
Loose ends affect a minority of cancer drivers. (a) Bar plot showing burden of amplifications via loose end by gene. Above, count of samples with at least 2 × *ploidy* copies of some part of the gene body and a loose end responsible for the gain, colored by type of repeat content identified at the loose end. Below, fraction of all gains of the gene caused by a loose end. (b) Bar plot showing burden of deletions via loose end by gene. Above, count of samples with fewer than 2 complete copies of the gene and a loose end responsible for the loss, colored by type of repeat content identified at the loose end. Below, fraction of all losses of the gene caused by a loose end. (c) Alluvial plot showing types of unmappable repeats identified at the seed and mate side of loose end rearrangements leading to the amplification of an oncogene. (d) Alluvial plot showing types of unmappable repeats identified at the seed and mate side of loose end rearrangements leading to the loss of a tumor suppressor. (e-g) Examples of loose ends impacting driver genes. Top, JaBbA graph representation of the sample showing loose end(s) responsible for gain or loss. Bottom, normalized binned read depth data, showing corresponding coverage depth changes. CN, copy number. RD, read depth. (e) Example gain of ERBB2 in esophageal carcinoma sample DO234446, with two loose ends mated to centromeric / pericentromeric sequence. (f) Example heterozygous loss of RB1 in TCGA glioblastoma multiforme sample 0221, with one loose end mated to centromeric / pericentromeric sequence. (g) Example homozygous loss of CDKN2A in TCGA lung adenocarcinoma sample 5899, with two loose ends mated to centromeric / pericentromeric sequence.

Among recurrently deleted cancer drivers, *FHIT* and *CDKN2A* had the highest burden of associated loose ends (50 and 44 instances respectively). *SMAD4* and *RB1* had the highest fraction of losses attributable to loose ends (10.2% and 8.7% of the total losses respectively) (**Fig. 7b**). This included examples of pairs of loose ends at *FHIT* loci that already harbored one or more deletion junctions classified as a simple deletion. Since *FHIT* has been previously shown to be a hotspot of rigma (a pattern of stepwise losses and clustered deletions at late-replicating fragile sites^39^), it is likely that the resolution of these additional loose ends would result in re-classification of these loci as rigma.

We next asked whether specific mutational mechanisms may generate loose ends at cancer genes. Cancer genes were significantly depleted in Type 2 and mystery loose ends relative to non-cancer genes, even after correcting for total gene breakpoint burden (*P* = 1.43 × 10^−7^, OR = 0.57), suggesting that they are less prone to NAHR. Type 1 loose ends within cancer genes showed a broad spectrum of mated repeat categories that largely reflected their genome-wide distribution. The most significant association was with centromeric or peri-centromeric sequence on the mate side (*P* = 0.027, OR = 1.48) relative to non-cancer gene-associated Type 1 loose ends (**Fig. 7c-d**). For example, an unclassified ten copy *ERBB2* amplification in an esophageal adenocarcinoma tumor sample (ICGC-D0234446) harbored a pair of Type 1 loose ends at opposite sides of the ~300 kbp amplified region, each linked to peri-centromeric sequence (**Fig. 7e**). These results are consistent with the presence of a double minute harboring junctional centromeric sequence. We found additional examples of Type 1 loose ends with centromeric or peri-centromeric mated sequences associated with *RB1* and *CDKN2A* loss in a glioblastoma (TCGA-0221) and lung adenocarcinoma (TCGA-5899) respectively (**Fig. 7f-g**). These results implicate that genetic instability around specific classes of repeats may contribute to the evolution of driver gains and losses.

## Discussion

As cancer WGS efforts scale into the tens or hundreds of thousands of samples and long molecule genome profiling technologies mature, it is important to rigorously approach the limitations of short reads in assessing somatic structural variation. The current expectation in the field is that short read WGS is very incomplete in its detection of SVs,^1, 50–52^ a view that has been advanced by long molecule profiling studies of limited sample sets (<5) of cancer cell lines.^63, 65–67, 69, 70^ The added sensitivity attributed to long molecules in these studies has been largely limited to the detection of complex somatic insertion and deletion variants smaller than 10 kbp.^52, 53, 71, 72^ While these studies have also identified large-scale SVs missed by short reads,^91^ such examples have been mostly limited to constitutional structural variation. Our approach to map and classify loose ends in cancer genomes thus represents one of the first analytic frameworks to rigorously assess the burden of missing structural data in somatic short read WGS.

Contrary to prevailing views, we find that a very small fraction (~2.3%) of large-scale cancer SVs are missed by short reads. Furthermore, the sum of Type 2 and mystery loose ends in our analysis provides an upper bound (1.2%) for the contribution of NAHR to cancer genomic structural variation. This validates the previously untested hypothesis that homologous recombination, which requires >97% sequence identity across long (>100 bp) sequences, plays a minor role in somatic structural genomic evolution. Our results suggest that certain cancer types (BLCA, BRCA, STAD, COAD, LUSC) have a higher burden (1.2% - 2.6%) of Type 2 and mystery loose ends and may require long molecule profiling to comprehensively characterize SVs and determine the contribution of NAHR. Our findings implicating nucleosome remodeling genes (*INO80* and *ARID1A*) in cancer associated NAHR merit functional validation.

Several factors may cause our loose end burden to underestimate the fraction of SVs missed by short read WGS. Our approach assumes that repeat-driven SVs will generate a read depth change in the bins surrounding an unmappable junction. Fully balanced rearrangements (e.g. reciprocal translocations, inversions) that are missed will yield a flat read depth profile without loose ends. However, balanced rearrangements are rare in the mappable cancer genome, representing approximately 0.68% of total rearrangements; even reciprocal rearrangements become unbalanced due to subsequent copy number changes including aneuploidies that alter the dose of one but not the other derivative allele. Though it is certainly possible that smaller inversions and late reciprocal events may be more likely to be balanced, a high rate of such events in the unmappable genome (but not the mappable genome) would imply a very specific mutational process that we believe is *a priori* unlikely. As a result, we conclude that undetected balanced rearrangements will obscure a very small fraction of repeat-driven SVs.

Second, certain breakpoints that arise in larger genomic “blind spots” where low mappability also distorts the regional read depth signal (e.g. centromeres) and may obscure true loose ends. However many large-scale SVs will still cause a read depth change in mappable bases that flank the genomic blind spot, which will shift but not obscure the loose end. Indeed, many of our mystery loose ends that are placed in mappable genomic regions and fail to generate tumor-specific chimeric contigs (**Fig. 2e**) may be the result of such a shift. As an example, many arm level copy changes occur through centromeric junctions and are detected in our analyses as centromeric or peri-centromeric loose ends (**Extended Data Fig. 6d-f**). However, we will not detect duplications or deletions that are fully contained inside the centromere. More generally, junctions whose breakpoints are local to that unmappable region and thus fail to cause a copy change in the surrounding bins will not generate a loose end. Excluding the centromeres themselves, we find that only 2.2% of the remaining genome harbors a long (≥ 10 kbp width) stretch of unmappable sequence. These results suggest that such local repeat-driven SVs likely contribute to a minority of the total SV burden.

The presence of Type 1 loose ends indicates that many somatic SVs arise through the fusion of unmappable repetitive sequence to mappable high-complexity genome. These include a high fraction of Type 1 loose ends associated with unmappable LINE or SINE elements. The fractional prevalence of these events (0.3% of all long-range SVs) even when combined with mappable SVs (~26%) is approximately consistent with their fractional contribution to the human genome. The contribution of these events to all SVs appears consistent across tumor types. These results indicate that the majority of LINE and SINE-associated somatic SVs are the result of random DNA breakage rather than specific SV driven mutational processes.

GRTR+ loose ends nominate locations where cancer chromosomes may have acquired new physical “ends”. The enrichment of GRTR+ (but not CRTR+) loose ends in *TERT* low and/or *ATRX* / *DAXX* mutant tumors suggest that cancer neotelomeres are not the result of telomerase-mediated healing^110, 111^ but arise in the context of ALT. These candidate neotelomeres are distinct from the interstitial telomere insertions that have been previously shown to be enriched in *ATRX* / *DAXX* mutant ALT cancers.^112–114^

How might the ALT phenotype drive neotelomere formation at interstitial double-stranded breaks? ALT cells have high concentrations of extrachromosomal telomeric DNA.^115^ These include short linear double-stranded telomeric DNA that provide abundant substrates for NHEJ to cap DSBs with short telomeric sequences. While such a process is equally likely to yield G- or C-rich 3’ ends, G-rich ends may be preferentially extended through rolling circle amplification after annealing to C-circles, which are ALT-specific extra-chromosomal circular telomeric DNA with fully intact C-rich and partially intact G-rich strands. Sufficiently long GRTR+ (but not CRTR+) ends will be stabilized into neotelomeres through shelterin complex formation, which requires G-rich 3’ overhangs to create t-loops and suppress further NHEJ.^116^ We note that the majority of tumors with GRTR+ loose ends were *ATRX* / *DAXX* wild type and showed negligible *TERT* expression. An intriguing possibility is that the presence of GRTR+ loose ends may mark tumors that rely on ALT for telomere maintenance despite lacking canonical ALT pathway mutations.

Long molecule profiling can help resolve loose ends into junctions; however, loose ends represent a minority of all large-scale SVs in cancer samples. The low frequency of loose ends at clinically important gene fusion partners suggests that long molecule profiling will only incrementally improve SV detection over state-of-the-art short read WGS pipelines. Given these findings, what sort of transformative insight can long molecule profiling provide into cancer genome structure? As we and others have previously shown,^38, 39, 43, 71, 117–120^ the allelic deconvolution of complex rearrangements can help define novel SV patterns and uncover mutational mechanisms. We propose that this sort of multi-junction phasing and scaffolding, rather than the resolution of individual loose ends, may be the most profound contribution that long molecules can provide into our understanding of cancer genome structure.

## Data availability

10X Genomics linked-read sequencing data for cell lines A549, HCC1143, Hs-294T, NCI-H1963, NCI-H209, NCI-H661, NCI-H526, U2OS, and HCC1143BL and Bionano Genomics optical mapping data for cell lines NCI-H838 and OVCAR-3 will be deposited in a public repository by the time of publication.

## Code availability

Software used in this paper can be found in the following GitHub repositories:

- https://github.com/mskilab/loosends
- https://github.com/mskilab/JaBbA
- https://github.com/mskilab/gGnome

## ACKNOWLEDGEMENTS

M.I., J.M.B., X.Y, A.D., J.R., and H.T. are supported by M.I.’s Burroughs Wellcome Fund Career Award for Medical Scientists, Doris Duke Clinical Foundation Clinical Scientist Development Award, Melanoma Research Alliance Team Science Award, National Institutes of Health U24-CA15020, and Weill Cornell Medicine Department of Pathology and Laboratory Medicine startup funds. K.H. is supported by a NIH/NCI F31 Graduate Research Fellowship (F31-CA232465). J.M.B, M.I., and T.d.L. are supported by Starr Cancer Consortium Award I13-0019.

## AUTHOR CONTRIBUTIONS

These contributions follow the Contributor Roles Taxonomy guidelines: https://casrai.org/credit/. Conceptualization: J.B., X.Y., T.d.L., M.I.; Data curation: J.B, X.Y., K.H., A.D., J.R., M.I.; Formal analysis: J.B., M.I.; Funding acquisition: T.d.L., M.I.; Investigation: J.B., T.d.L., M.I.; Methodology: J.B., X.Y., M.I.; Project administration: M.I.; Resources: M.I.; Software: J.B., X.Y., M.I.; Supervision: M.I.; Validation: J.B, H.T., J.R., M.I.; Visualization: J.B., M.I.; Writing – original draft: J.B, M.I.; Writing – review & editing: all authors.

## Methods

### Obtained Data Sources

Cancer genome short read WGS data for 2319 tumors and cell lines used throughout this study were obtained for a previous study published in.^39^ 10X Genomics linked-read WGS for three cell lines (HCC1954, HCC1954BL, and NCI-H526), used to compare short read sequencing with long molecule technologies, were also previously obtained for the same study.

Additionally, for cell lines for which optical mapping and linked read libraries were generated, corresponding whole genome sequencing data were obtained. WGS data for cancer cell lines A549, Hs-294T, NCI-H1963, NCI-H209, NCI-H526, NCI-H661, NCI-H838, and Ovcar-3 were obtained from the Cancer Cell Line Encyclopedia (CCLE) (https://portals.broadinstitute.org/ccle),^121, 122^ with permissions granted by the proprietors of this dataset.

### Reference Sequence and Annotation Sources

Cytoband and repeat tracks were obtained from the UCSC Genome Browser database. The repeat sequence and poly-A databases from TraFiC (,^56^ github download, https://gitlab.com/mobilegenomesgroup/TraFiC/~/tree/multispecies/databases/hg19) were supplemented with ri-bosomal reference sequences and satellite reference sequences from RefSeq (alpha-satellite consensus sequence X07685.1, gamma X satellite sequence X87951.1, ribosomal complete sequence U13369.1, beta-satellite sequence M25749.1) to comprise the repeat reference against which contigs were aligned. 6251 viral sequences were also obtained from RefSeq v1.1 (ftp://ftp.ncbi.nlm.nih.gov/refseq/release/viral/).

### 10X linked-read whole genome sequencing

Twelve cell lines (cancer cell lines A549, Hs-294T, NCI-H1963, NCI-H209, NCI-H526, NCI-H661, NCI-H838, U2OS, HCC1954, HCC1143, and blood cell lines HCC1954BL and HCC1143BL) were subjected to 10X Chromium linked-read whole genome sequencing. 10X linked-read sequencing BAMs aligned to assembly GRCh37. HCC1954, HCC1954BL HCC1143, and HCC1143BL cell lines were obtained from 10X Genomics.

High molecular weight (HMW) genomic DNA (gDNA) was extracted using a Qiagen MagAttract HMW DNA Kit (Qiagen, Germany) according to the suggested protocol. Approximately 2 million fresh cells were lysed, HMW gDNA was captured by magnetic particles (Qiagen MagAttract Suspension G), and then the magnetic particles with HMW gDNA was washed in wash buffer and eluted in EB Buffer (10 mM Tris-HCl, pH 8.5). The HMW gDNA had a mode length of 50 kbp and max length 200 kbp, as estimated on a separate 75V pulse-field gel electrophoresis (BluePippin 5-430kbp protocol).

10X sequencing library preparation was performed using a Chromium Genome Library Kit v2 (Lot 152527, 10X Genomics) following the Chromium Genome Reagent Kits v2 User Guide. 1 ng of extracted HMW gDNA was used to prepare a 150 bp paired-end library, with an average fragment length of 625 bp (ranging from 300 to 2000 bp, measured with the Bioanalyzer High Sensitivity DNA Kit, Agilent). The prepared library was sequenced on an Illumina NovaSeq 6000 Sequencing System with S4 flow cells, to an average read depth of about 33X, resulting in approximately 173X physical coverage for NCI-H838. All the 10X linked reads were aligned with Long Ranger (v2.1.3, 10X Genomics).

### DNA isolation for optical mapping

Ultra high molecular weight DNA extraction was performed on cell lines NCI-H526, NCI-H838, and Ovcar-3 using a Bionano SP Blood & Cell DNA Isolation Kit catalog #80030, Bionano Genomics, San Diego), according to the Bionano Prep SP Frozen Cell Pellet DNA Isolation Protocol (document #30268, revision B). 1.5 million frozen cells in cryopreservation medium were thawed in a 37°C water bath, centrifuged for 2 minutes at 2000xg, washed, and re-suspended in DNA Stabilizing Buffer, made with 2% v/v DNA Stabilizer (PN 20397, Bionano Genomics) in Cell Buffer (PN 20374, Bionano Genomics). Cells were treated with proteinase K (#158920, Qiagen, Germany) and RNase A (#158924, Qiagen) in the presence of detergents and salts. DNA was bound to a silica disk, washed, eluted, and homogenized via 1 hour of end-over-end rotation at 15 rpm. The ultra-high molecular weight DNA was allowed to rest overnight at room temperature before fluorescent labeling.

### Bionano fluorescent DNA labeling and imaging

Ultra-high molecular weight DNA was fluorescently labeled at the motif CTTAAG with the enzyme DLE-1 and counter-stained using a Bionano Prep™ DNA Labeling Kit – DLS (catalog #8005, Bionano Genomics, San Diego) according to the Bionano Prep Direct Label and Stain (DLS) Protocol (document #30206, revision F). DNA imaging and generation of single-molecule reads was performed on a second Generation Saphyr System with Saphyr chips (#60325 and #20366, Bionano Genomics) running Instrument Control Software version 4.7.18339.1. At least 400X coverage was generated.

### JaBbA loose end mathematical formulation

As defined in,^39^ the JaBbA algorithm infers junction-balanced genome graphs (JBGGs) from junctions and breakends obtained through the analysis of cancer whole genome sequences (WGS) by fitting integer vertex and edge weights to high-density WGS read depth data through the solution of a mixed integer quadratic program (MIQP). A genome graph is a directed graph *G* = (*V, E, ψ, φ*) whose vertices *v* ∈ *V* represent strands of DNA sequences, and whose edges *e* = (*v*_1_*, v*_2_) ∈ *E*(*G*) represent genomic adjacencies (i.e. 3-5’ phosphodiester bonds) joining two DNA sequences, where *v*_1_*, v*_2_ ∈ *V* (*G*). Loose ends are incorporated at sites of disagreement between fitted vertex weights and their corresponding edge weights, as described below.

#### Junction-balanced genome graph

We define a mapping *κ*: {*V_I_* ⋃ *E* → ℕ of non-negative integer copy number (CN) to vertices and edges of *G*, where *κ*(*v*)*, v* ∈ *V_I_* and *κ*(*e*)*, e* ∈ *E* represent the CN of vertex *v* and edge *e*, respectively. The principle of *junction balance* constrains the CN of every vertex to be equal to the sum of its incoming edges and the sum of its outgoing edges. Formally, the junction balance constraint is stated as follows:

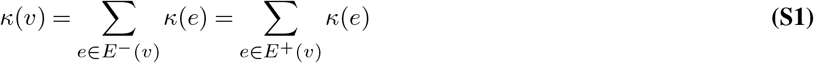

In addition we require the CN *κ* to obey *skew-symmetry*, which means that every vertex must have the same copy number as its reverse complement.

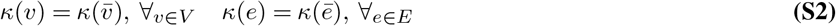

We call the combination (*G, κ*) for which *κ* satisfies Eqs. S1-S2 a *junction-balanced genome graph* (JBGG).

#### Inferring Junction Balanced Genome Graphs

We infer JBGGs from a genome graph *G* and binned, normalized, and purity / ploidy-transformed read depth data *x* ∈ ℝ^*n*^ across *n* genomic bins (see below for read depth transformation details) through the solution of a mixed integer quadratic program (MIQP), which assigns an integer CN *κ*: *V_I_* ⋃ *E* → ℕ to the vertices and edges of *G*. The genome graph *G* is generated, as above, from a set of breakends 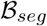 obtained from a preliminary segmentation of genome-wide read depth (i.e. via segmentation software such as CBS) and a set of junctions 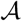 (i.e. from a junction caller such as `SvABA` or DELLY).

Each vertex *v* ∈ *V_I_* (*G*) is associated with a partition of bins *J* (*v*) ⊆ {1*,…, n*} (based on genomic coordinate overlap) and a mean bin value 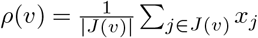. We model each bin subset 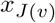 as an i.i.d. sample from a Gaussian distribution with standard deviation *σ*(*v*) and mean *κ*(*v*). The log likelihood is

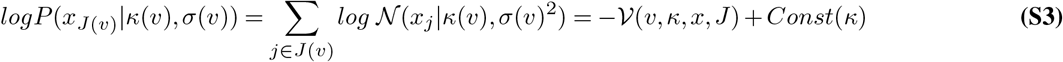

where 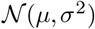 is the Gaussian probability density function with mean *μ* and variance *σ*^2^ and 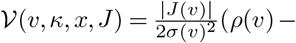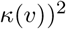 is the *read depth residual* of vertex *v*. The variance *σ*^2^(*v*) is a *κ*-independent parameter that models read depth noise and is computed directly from the data. The simplest noise model is a constant, where this parameter is set to the genome-wide sample variance of the read depth around each vertex mean: 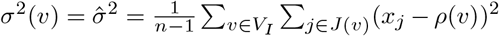. In practice, we apply a vertex-specific variance estimate *σ*^2^(*v*) to account for heteroscedasticity in the read depth data (see “JaBbA model fitting” section below).

Given this model, the joint log-likelihood of the read depth data *x* across the graph given copy number assignment *κ* is

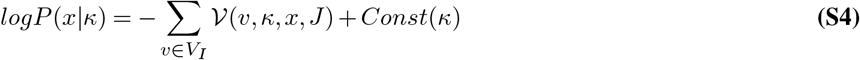

We also refer 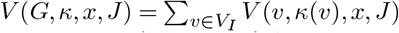 as the *read depth residual* of the JBGG (*G, κ*) relative to data *x*.

The satisfaction of junction balance and skew-symmetry constraints in Eq. S1-S2 may place nonzero copy number at one or more loose end edges. Each loose end in the input graph represents a slack variable that allows the junction balance constraint to be relaxed at specific internal vertices, allowing the data to be fit even when junctions are missing from the input (e.g. due to low mappability, sequencing depth, or purity). Only loose ends that are given nonzero CN, are considered to be “used” in the final graph. To penalize solutions that require the use of many loose ends, we add an exponential prior with decay parameter *λ* on the loose end CN in (*G, κ*), which makes models with many missing junctions unlikely. This prior has log likelihood

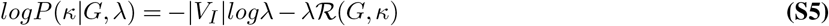

Where

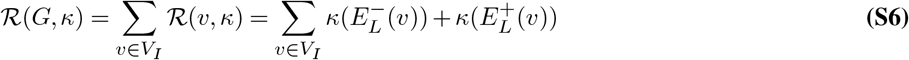

is a *complexity penalty*. Adding the log likelihood in Eq. S4 to the prior in Eq. S5 yields a penalized log likelihood for the data with regularization parameter *λ*. Under this model, the maximum a posteriori probability (MAP) estimate of *κ* will minimize the function

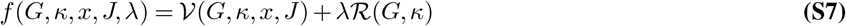

which combines the quadratic read depth residual 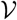 and *ℓ*_1_-norm complexity penalty 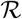 into a single quadratic objective. In practice, we apply models that penalize the number of loose ends with nonzero copy number, i.e. applying an *ℓ*_0_-norm penalty 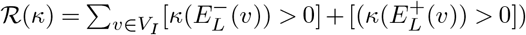. We use *f* to define a MIQP, which we solve to infer a MAP estimate for *κ* given data *x* and genome graph *G*:

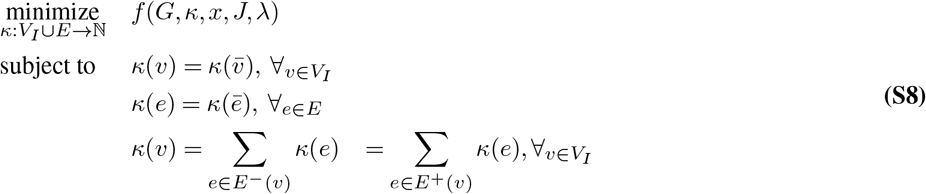

The resulting MAP estimate 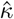 defines the JBGG 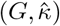 which is outputted and returned to the user.

### Representing mappability of hg19 coordinates

At every position in the GRCh37/hg19 reference genome, the 150mer of reference sequence originating from that position was extracted and written to fastq, with a string representing the source coordinate as qname. This fastq was aligned to the GRCh37/hg19 reference using Burrows-Wheeler aligner software^123^ (`bwa mem`, v.0.7.10-r789). The alignments were then parsed to extract the qname (representing the base of origin of the 150mer sequence) and the alignment MAPQ. The MAPQ was assigned to the base of origin of the sequence, regardless of the alignment position. Every base in the reference genome is therefore annotated with a MAPQ score for a 150bp read originating from that base. Reference coordinates assigned MAPQ=60 (the maximum for `bwa mem` v.0.7.10-r789) are considered mappable, and all others unmappable. The same process was repeated for 101mer sequences.

### Mappability of RepeatMasker annotations

Each base in the reference genome was individually annotated as: (1) mappable or unmappable for a given read length and (2) whether that base overlapped any RepeatMasker annotation. Centromeric coordinates were defined as all bases within the last p-arm and first q-arm cytobands, excluding bases that already had a RepeatMasker annotation.

### Loose end quality filters

Prior to running JaBbA, normalized read depth data has been preprocessed as described in.^39^ Genome-wide 200 bp bins are annotated with GC and mappability corrected read depth profiles for the tumor and paired normal, as well as the tumor / normal ratio. For each genome graph, every non-terminal fitted loose end was evaluated. A sample-specific value *β*, which represents the magnitude of coverage change corresponding to one copy state change throughout the sample, is calculated based on **Equation 11** from.^39^ In essence, the definition of *β* is as follows:

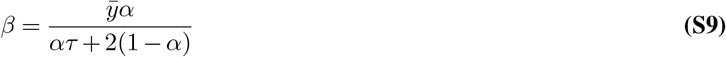

where *ȳ* represents the average normalized read depth genome-wide, *α* is the sample purity, and *τ* is the sample ploidy. Loose ends within an unconverged subgraph of the JaBbA model were filtered out of our dataset.

The coordinates for the fused and unfused sides of each loose end were defined by the coordinates of nodes flanking the loose ends, up to a maximum of 100 kbp from the loose breakpoint. Read depth data for bins falling within the fused and unfused sides of every loose end is further analyzed. Bins with ratio values in the bottom or top 5th percentile for every side of every loose end are excluded.

Autocorrelation of ratio values along reference coordinates were quantified to identify false positive CNA calls driven by misleading coverage noise. For every fused or unfused side of every loose end, the autocorrelation (acf function from the stats package within R with argument “type = correlation”) of the ratio value is calculated at every lag value between 5 kbp and 100 kbp (or the length of that side, if shorter than 100 kbp), and the sum of every autocorrelation estimate squared is assigned to the corresponding loose end side. The loose end is assigned the maximum value of the corresponding fused and unfused sides. An empirically chosen maximum threshold of 2 was used to define an acceptable level of autocorrelation. Loose ends assigned an autocorrelation value ≥ 2 were filtered out of the dataset.

A generalized linear model is then fitted to the tumor and normal coverage values (glm function from the stats package within R with argument “family = gaussian” to score coverage values as a function of their start position, sample, and side (fused or unfused)). The residual between the value predicted by the model fit and actual coverage value is calculated for each input coverage value, and a Kolmogorov–Smirnov (KS) test (ks.test function from the stats package within R) was performed to compare the distributions of residual values for tumor sample coverage on the fused versus unfused sides of the loose end, and the loose end is annotated with the resulting P-value. Loose ends with *P* ≥ 0.01 were filtered out of the dataset.

Additional KS tests are performed to compare tumor coverage values on the fused versus unfused sides, and normal coverage values on the fused versus unfused sides. The loose end is annotated with both resulting P-values.

The magnitude of coverage ratio change between the fused and unfused sides was calculated by taking the difference of the ratios of median tumor coverage value / median normal coverage value on the fused versus unfused sides of the loose end. The magnitude of change from the fused to unfused sides of the loose end is also calculated within the tumor coverage values (median tumor coverage value on the fused side - median tumor coverage value on the unfused side) and within the normal coverage values. Loose ends with a gain of magnitude in either the tumor coverage or tumor / normal ratio below a threshold of 0.6 × *β* were filtered out of the dataset. Loose ends with a gain of magnitude in the normal coverage above the same threshold were also filtered out of the dataset. Loose ends with a gain of magnitude in the tumor / normal ratio less than the standard deviation of coverage ratio values within the fused or unfused side of the loose end were also filtered out of the dataset.

The entire dataset of loose ends were loaded in aggregate, and the P-values from the three KS tests were adjusted with Bonferroni correction to nominate statistically significant associations (`p.adjust` within R). Loose ends included in the final dataset of 26751 were those with a Bonferroni < 0.05 for the KS test of fitted residuals, Bonferroni > 0.05 for the KS test of normal coverage values, and Bonferroni < 0.01 for the KS test of tumor coverage values.

### Identification, realignment, and annotation of loose reads

At every loose end, reads aligned within 5 kbp from the breakpoint in the corresponding bam file are loaded into R using `scanBam` from Rsamtools. The metadata fields `RNEXT` and `PNEXT` and tag `SA` (which provides information about split alignments of a single read) are parsed to identify coordinates for all mates and split alignments of the reads within the 5 kbp window. The bam is then parsed again using `scanBam` at these new coordinates, to load all mate sequences into R. Reads loaded in the second round that do not share a qname with reads loaded in the first round are discarded. The sequences of reads that were aligned to the negative strand of the reference genome are reverse complemented, such that all read sequences are in their original reading frames. The read sequences are realigned single-end to the GRCh37/hg19 reference genome within R using a `BWA` object from package `RSeqLib`. Single end alignment allows each individual read to have its own MAPQ, as paired end alignment will assign a single MAPQ to the entire pair. Following realignment, loose read pairs are identified by annotating qnames for which one read has been assigned a MAPQ = 60 and the other has a MAPQ = 0. The reads in these pairs with MAPQ = 60 are loose reads, while the reads with MAPQ = 0 are loose mates. The alignment positions of loose reads only, not loose mates, are assessed for proximity to loose breakpoints.

Overlaps were found between the alignments of loose mates and RepeatMasker annotations. Loose read pairs were than annotated by the type of repeat (if any) overlapping the loose mate alignment.

### Assembling local read pairs

Strand-specific tumor and normal contigs are assembled from a seed region within 1 kbp from the loose breakend. The seed region is divided into 200 bp bins. For a given bin, the assembly process is performed four times: once each for the positive and negative strand in the tumor and normal samples. All read sequences begin in their original reading frame by converting the sequences of reads aligned to the negative reference strand to their reverse complements. Reads from the appropriate sample and strand are incorporated if their alignment overlaps any bases belonging to the bin. Their sequence is kept in the original reading frame orientation. The mate pairs of these reads are also incorporated, with their sequences reverse complemented from the original reading frame orientation. If both mates belonging to a read pair align to the bin on the same strand, one is arbitrarily chosen to reverse complement. If there are at least six total reads, all of the read sequences are then used to construct a `Fermi` object (from package `RSeqLib`, argument “assemble = TRUE”) to assemble. If any contig sequences are identified, they are used to construct a `BWA` object. Because the input sequences to `Fermi` represented a single strand, all input sequences are expected to align to the same strand of resulting contigs. Any contig sequence resulting in more alignments to its negative strand than positive is replaced by its reverse complement. Every contig is annotated by the number of reads aligned to it with MAPQ=60 when aligned to GRCh37/hg19.

### Aligning loose read contigs

Assembled contigs are aligned to reference genomes in two stages. First, the contig sequences are aligned to the GRCh37/hg19 human reference. They are also aligned to a repetitive element database containing consensus sequences for LINE-1, Alu, SVA, ERVK, polyA, ribosomal, and satellite elements. The cigar strings for all alignments are then parsed to identify any substrings of contig sequences with no reference alignments. Contig sequences with unaligned substrings are subsequenctly aligned to a combined reference genome which includes the GRCh37/hg19 reference and 6251 viral genome sequences. Any resulting alignments to viral genomes are only appended to the initial stage alignments if they include an alignment for previously unmapped substring sequence, again identified by parsing the alignment cigar string. Alignments to GRCh37/hg19 are checked for overlaps with a custom track of unmappable repeats. This track starts with RepeatMasker annotations that contain ≥ 50% unmappable bases, calculated as previously described. All rare repeat classes (any annotation not included in LINE, SINE, LTR, Simple repeat, Satellite, and Low complexity) are classified as “Other”. Centromeric regions are identified by first generating 3 Mbp sliding windows genome-wide every 100 kbp. Each sliding window is annotated by the fraction of bases within that window that are unmappable. Windows that are at least 50% unmappable are collapsed via the `reduce` function from R package `GenomicRanges`. Reduced windows that overlap at least one base of a centromeric cytoband are joined with cytobands staining for either acen or gvar, and the total union of these three groups are classified as centromeric / pericentromeric coordinates. The first and last cytoband of each chromosome are classified as telomeric / subtelomeric coordinates. The unmappable repeat track is a union of the unmappable RepeatMasker annotations, centromeric coordinates, and telomeric coordinates. Any overlaps between these annotations and contig alignments to GRCh37/hg19 are appended to the contig alignments with an annotation indicating the type of repeat found at the overlap.

### Categorization schema for loose end taxonomy

All tumor-originating contigs assembled around a given loose end are input to the categorization schema. Contigs assembled from a seed bin not within 1 kbp of the loose end on the loose end supporting strand are filtered out. The seed locus for each contig is assessed for overlaps with alignments of that contig, with a pad of 750 bp representing possible read insert lengths that could separate concordant alignments. Any contig alignment overlapping its own seed within 750 bp on the same strand represents the “seed” portion of the contig. If any contigs contain seed sequence, and all contig seed alignments align to GRCh37/hg19 with MAPQ=60, the loose end has a mappable seed. Alignments to GRCh37/hg19 with a MAPQ < 60 are annotated as repeats with repeat type “unmappable-NOS” (NOS = not otherwise specified).

The contig alignments are then converted into contig coordinates by parsing the cigar strings of the alignments. Alignments are supplemented at this stage by parsing the sequences for matches to a telomeric 18mer, using a `PDict` object from package `Biostrings` as described in more detail below. Any matches to the 18mer sequence are converted into contig coordinate ranges representing the matching bases and annotated as “G telomeric” or “C telomeric” depending on which orientation of the telomere motifs had a match. If viral alignments share at least 90% of their bases with telomeric sequence matches, they are removed from the contig alignments. If ribosomal alignments share at least 90% of their bases with polyA alignments, they are removed from the contig alignments. If coordinate-based telomeric alignments (i.e. alignments to GRCh37/hg19 that overlapped telomeric / subtelomeric regions as defined by the unmappable repeats track) overlap telomeric sequence matches, they are removed from the contig alignments to prioritize the more specific sequence match annotation.

All contig alignments in contig coordinates overlapping an alignment that was annotated as “seed” are treated as features of the seed region. All contig alignments downstream of the seed region in contig coordinates are treated as features of the mate. If any repeats are present on the seed side, they are annotated as “seed repeats”. If any repeats are present on the mate side, they are annotated as “mate repeats”. If no contig has at least 10 MAPQ=60 reads supporting it, an “unmappable-NOS” annotation is added to the seed repeats. Contigs with alignments to the seed region within the last (3’) 20 bases of the contig sequence are filtered out, after collecting annotations for any seed repeats that may be present.

If the seed sequence has an alignment to GRCh37/hg19 with MAPQ=60, the seed is mappable. If the mate sequence has an alignment to GRCh37/hg19 with MAPQ=60, the mate is mappable. If the seed and mate are both mappable, and the seed and mate alignments for all contigs represent a single pair of loci joined in the same orientation, the loose end will be classified as Type 0. If the seed and mate both have MAPQ=60 alignments, but do not represent a single consensus junction, the mate is reclassified as unmappable. If no mate repeats are present, a “complex” annotation is added to the mate.

If no contigs were assembled or remain after filtering, the loose end coordinate is compared to the unmappable repeat track described above. An overlap between the unmappable repeats and the loose end will be categorized as an unmappable seed; the loose end is otherwise a mystery. For loose ends in the mystery category, the asssembly, alignment, and categorization process is repeated beginning from three 1 kbp sliding windows every 500 bp within the 1 kbp window surrounding the loose end to use as seed bins.

### Associating contigs with repeat types

All seed and mate repeats from all contigs for a given loose end that remain after filtering are analyzed to choose one predominant repeat type for each side of a loose end rearrangement. The options for a seed or mate repeat annotation are centromeric / pericentromeric, satellite, LINE, low complexity, LTR, poly-A, ribosomal, simple repeat, SINE, telomeric / subtelomeric, G telomere, C telomere, unplaced hg19 contigs, unaligned sequence, viral, other repeats, and unmappable-NOS. “Unplaced hg19 contigs” refer to the contigs in the GRCh37/hg19 reference assembly which have not been incorporated into one of the 24 chromosome assemblies. “Unaligned sequence” indicates that a significant portion (at least 20 bp) of the rearrangement contig did not align to any reference or repetitive sequence database. After filtering certain overlapping repeats described above, the longest repeat alignment is chosen as the predominant repeat type, with the exception of unmappable-NOS, which will only be chosen as the predominant repeat type if no other repeat types are present.

In downstream processing, the centromeric / pericentromeric and satellite categories have been merged, as have the telomeric / subtelomeric, G telomere, and C telomere categories. The original annotation is retained, and only G telomere matches are used to identify neo-telomeres.

### Type 0 rescue by discordant read pairs

Discordant read pair alignments surrounding the loose ends were additionally analyzed to identify Type 0 junctions and Type 1 loose ends with complex rearrangements. Beginning again from 200 bp bins within 1 kbp from the loose end, reads from the tumor sample aligned to the loose end-supporting strand and overlapping the bin are pooled along with their mates. Same-stranded overlaps of the alignments of at least 10 reads with MAPQ=60 are identified, and identified as either “seed” (overlapping the strand-specific 200 bp bin) or “mate” (sharing no strand-specific overlap with the bin). If all bins harboring at least ten discordant read pairs identify the same pair of seed and mate loci, that loose end will be annotated as a missed junction. If multiple mate loci are identified, the loose end will be annotated as Type 1.

### Quantifying sequence entropy of assembled and unassembled reads

To assess whether low complexity read sequences were contributing to mystery loose end designations, the entropy of all tumor reads used as inputs to the `Fermi` assembler on the loose end supporting strand were quantified. The strand-specific contigs were used to instantiate a `BWA` object, and the seed-aligned reads and their mates were aligned to it. Individual reads (not pairs) that align to the contig with at least 90% of their sequence included in the alignment are “assembled”. All other reads are “unassembled”. All unassembled reads and a random sample of an equivalent number of assembled reads are divided into sequence 3mers. The count of instances of every possible sequence 3mer present in each single read is input to the `entropy.empirical` function from the R package `entropy`. Each read is assigned a single entropy score.

### Recategorizing loose ends based on long molecule SV calls

Loose ends “rescued” by long molecule sequencing could be recategorized into Type 0, Type 1, or Type 2 based on the original classification of the seed locus and characteristics of the reference sequence at the mated side of the long molecule SV. Because the resolution of OM is much lower than LR, only LR-rescued loose ends were recategorized. Each side of the rescuing SV was separately determined to be mappable or unmappable, and then the SV (and rescued loose end) were categorized as Type 0, Type 1, or Type 2 based on whether both sides, one side, or neither side were mappable, respectively. Each loose end is recategorized separately, even if the LR SV indicates a junction between two short read loose ends. For each loose end, the mappability defined by short reads is assigned to the seed side of the LR junction. If the mate side of the LR junction does not fall within 3 kbp of a short read CNA, the mate is automatically considered unmappable. Next, a 100 bp window extending from the mated junction breakpoint into the fused sequence was considered. If any bases within this 100 bp window had MAPQ < 60, the site is considered unmappable. Additionally, the fraction of bases with MAPQ < 60 were counted within a 3 kbp pad around the 100 bp window. If the fraction was ≥ 0.2, the site is considered unmappable.

### Associations between loose ends and complex SV patterns

Complex SV patterns were identified on JaBbA graphs using the `events` function of R package `gGnome`. Every pattern has a corresponding genomic footprint. Only samples with at least 10 total breakpoints (associated with junctions or loose ends) were considered. For every instance of an event, sample-specific loose ends falling within a 100 kbp pad around the footprint were counted for 41 tumors: the tumor the event came from, and a random sample of 40 other tumors in the cohort that also contain at least one instance of the event type. For each event type, associations between the event type and loose ends were quantified by a generalized linear model (`glm` function from the `stats` package within R with argument `family = binomial(link = ‘logit’)`). The GLM was fitted to a logical indicator of whether any sample-specific loose ends were found within the footprint coordinates as a response to a logical variable indicating whether the sample in question contained the event at that footprint, the total count of junction break points in that sample on that chromosome, and the width of the event footprint.

Associations between G-rich telomeric loose ends and complex SV patterns were calculated with a similar formulation, with two modifications. All G-rich telomeric loose ends sharing a chromosome with the event footprint were counted (regardless of alignment within a 100 kbp pad around the footprint). An additional logical covariate was used describing whether the sample contained any instances of G-rich or C-rich telomeric loose ends.

### SV calling from optical mapping data

The rare variant pipeline (Bionano `Solve` version 3.X) was used on the optical mapping dataset fluorescent labeled molecule images to optically map and detect junctions. The junction-calling pipeline consists of three major steps: initial molecule alignment and clustering of junctions, consensus generation by molecule extension refinement, and final junction calling.

Using `RefAligner` (version 9248,,^124^,^125,126^), molecules in an input BNX file are aligned directly to the reference hg19 `(-T 1E-7, -A 5, -L 40, -S 0.1 -sv 3 -MultiMatches 5)`. Significant internal alignment gaps `(-outlier 1E-2)` and end alignment gaps `(-endoutlier 1.1E-2)` may indicate the presence of junctions in the sample. To identify low allelic frequency junctions, the pipeline requires only a default minimum of three molecules calling the same junction. Molecules are determined to confirm the same insertion (> 5 kbp in size), deletion (> 5 kbp), duplication (> 25 kbp) or inversion (> 50 kbp) if the inferred variant positions overlap and their inferred variant sizes are similar (default of 20% size similarity). To confirm the same translocation, the inferred translocations must be in the same orientation and their breakends in close proximity (within a distance of 35 kbp).

A consensus is built using clusters of molecules that identified the same junction. The purpose of the consensus step is to verify that those junction-supporting molecules truly agree and can form a consensus that represents the variant allele. In this step, the loci on the reference assembly 150 kbp flanking the inferred junction are extracted, and the molecules that supported the junctions are aligned to each of the two junction-flanking reference fragments. The pipeline attempts to reconstruct the junction allele by using the molecules to extend into the junction region `(-T 1E-7 -A 5 -L 40.0 -S 0.1 -extend -maxExtend 250.0)`. If the molecules come from the same variant, they would have similar label patterns and form a consensus map that represents the variant allele. Note that for each junction, the same extension procedure is performed twice: one extension from the fragment left of the junction and one extension from the right.

Finally, the new local consensus maps are realigned to the reference to check if the same initial junction calls are made `(-T 1E-7 -A 5 -L 40 -S 0.1 -sv 3 -MultiMatches 5 -svAlignConfByInterval 0.7)`. The pipeline will only report junctions that are confirmed in the final junction calling step. Note that potentially two consensus maps could form for each junction, but only one is kept in the end.

breakpoints of junction calls identified by the Bionano `Solve` rare variant pipeline “rescue” a loose end from the same sample if they fall within 10 kbp of the loose end coordinate on the same strand.

### SV calling from linked read data

Bam files generated by the `longranger lariat` alignment pipeline contain barcode information as a tag assigned to each read entry. These bams were used as input to three linked read SV calling algorithms: `GROC-SVs, NAIBR`, and `LinkedSV`.

breakpoints of junction calls identified by the union of all three linked read SV callers “rescue” a loose end from the same sample if they fall within 3 kbp of the loose end coordinate on the same strand.

### Identifying telomeric 18mer matches in hg19 reference sequence

Telomeric motifs included in this analysis are TTAGGG, TCAGGG, TGAGGG, and TTGGGG, and their reverse complements. Every possible 18mer substring of a strand-specific combination of any of these motifs was loaded into two strand-specific `PDict` objects from package `Biostrings`, resulting in a total of 832 unique 18mers (416 for each strand orientation). The 150mer reference bam file was used to find sites of telomeric 18mer matches in the reference genome. Reads were loaded into R using the base `pipe` function to stream the output of `samtools view` from the command line. The read sequences were instantiated as a `DNAStringSet` object from package `Biostrings`. Using the function `vwhichPDict` from the same package, a logical value is assigned to every read indicating whether it contained an exact match for any of the 18mers, for both strand orientations. For all reads with positive matches to 18mers (from either strand), the original source coordinate of the read was extracted from the qname. The entire 150 bp region for each read is considered a positive match for telomeric 18mer content, and used to quantify the fraction of the genome with positive matches. The first and last cytobands of every chromosome were used to represent telomeric and subtelomeric regions, and overlaps with these cytobands were used to quantify the fraction of 18mer reference matches falling within or outside of telomeric regions.

### Identifying telomeric reads genome-wide

To generate aggregate tracks of telomeric read peaks in real samples, a similar approach was used as the previous, with an additional intermediate step. After all read sequences were parsed for matches to the telomeric 18mers, the entire bam file was streamed using `pipe` for `samtools view` again, to extract all reads with the same qname as a read with a telomeric 18mer match. These reads and their mates were realigned single end to GRCh37/hg19 using a `BWA` object in R from package `RSeqLib`. Individual reads were aggregated if their sequence did not contain an 18mer match but the sequence of their mate did. The `coverage` function from `GenomicRanges` was used to identify peaks of at least 5 G-rich mated reads aligned to the same coordinates in a single sample. All peaks were compared against the breakpoints of nodes from the corresponding JaBbA graphs to determine whether they fell within 1 kbp of a loose end (“At loose end” in Fig. 5), within 1 kbp of the end of a node that is not loose (filtered out of analysis) or not near a node breakpoint (called “Flat CN”). The peak coordinates and strands were then used as reference positions around which the alignments of all telomere-mated reads were aggregated.

### Identifying loose ends responsible for oncogene gains

Oncogenes included in this analysis were COSMIC Cancer Gene Census (CGC,^127^) version 86 genes that intersect with the pan-cancer GISTIC amplification peaks in Zack et al, 2013.^109^ Loose-end driven oncogene gains were counted in this analysis if any part of the gene body of the oncogene is amplified to at least ≥ 2 × *ploidy* copies and a loose end is responsible for at least ploidy copies. All overlaps between oncogene bodies and nodes present at ≥ 2 × *ploidy* copies were first identified. Subgraphs around these amplifications were then expanded one node at a time until a loose end was reached. As described in,^39^ potential haplotypes *H* are enumerated by identifying all traversals through the local subgraph *G* from source to sink vertices. A modified version of Equation 13 from that paper, defined as follows, is used to assign a copy number to each allelic haplotype subject to the constraints of the local vertex and edge copy numbers.

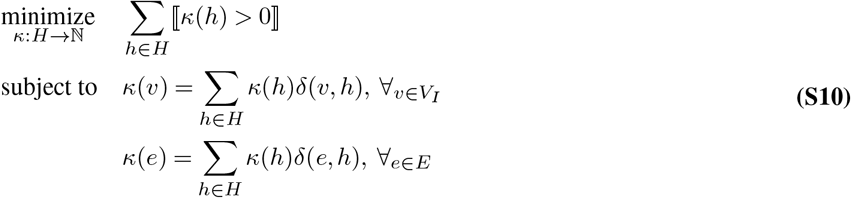

This mixed integer problem (MIP) is solved using `Rcplex` with argument “n = 200” to generate up to 200 possible allelic solutions to the subgraph. Each allele ‘h’ is annotated for whether it contains both the node containing the amplified oncogene body and the loose end. All of the generated solutions are parsed to extract how many total copies of alleles *h* ∈ *H* containing both the oncogene and loose end are present. The minimum value of these total copies from a single solution is taken as the minimum number of copies of the oncogene directly attributable to the loose end. If this value is at least ploidy, this is counted as an instance of a loose end causing an oncogene gain.

### Identifying loose ends responsible for tumor suppressor losses

Tumor suppressor genes included in this analysis were COSMIC Cancer Gene Census genes that intersect with the pan-cancer GISTIC deletion peaks in Zack et al, 2013.^109^ Loose- end driven tumor suppressor losses were counted in this analysis if the any part of the gene body of the tumor suppressor is present at < 2 copies and falls on the unfused side of a loose end. All overlaps between tumor suppressor gene bodies and nodes present at < 2 copies were first identified. Subgraphs around these depletions were then expanded one node at a time until nodes at ≥ 2 copies were reached in both directions. If a loose end is present on a terminal node of the subgraph on the strand facing towards the tumor suppressor body, this loose end is considered responsible for the tumor suppressor loss.

### Enrichment of centromeric or peri-centromeric sequences in Type 1 loose ends impacting driver genes

Type 1 loose ends were used to quantify cancer gene associations. Every Type 1 loose end (6,530 total) was annotated with a TRUE/FALSE indicator for whether that gene was responsible for the loss or gain of a cancer gene, and another TRUE/FALSE indicator for whether centromeric or peri-centromeric sequences (including alpha-satellite matches) were found in the mated sequence. Significance was determined using Fisher’s exact test.

## Extended data figures

**Extended data Fig. 1.**
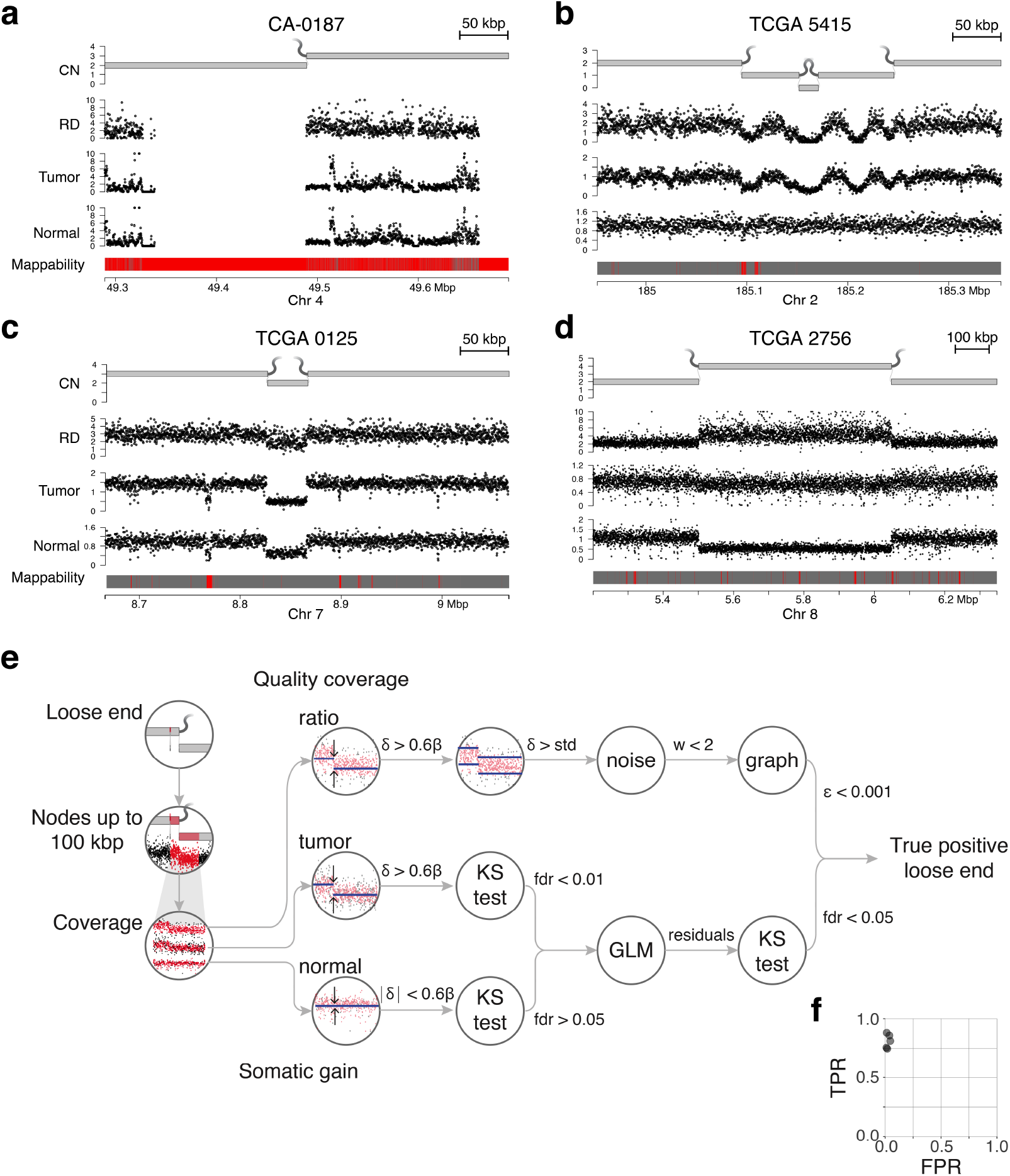
Loose end cohort. (a-d) Representative examples of filtered loose ends. Each from bottom: mappable regions of the genome (based on 150 bp reads) in gray, unmappable in red. Normalized binned read depth in paired normal sample. Normalized binned read depth in tumor sample. Transformed ratio of tumor/normal coverage depth. JaBbA graph representation of sample. CN, copy number. RD, read depth. (a) A single loose end in Cancer Alliance bladder urothelial carcinoma sample CA-0187. This loose end falls within the centromere of chromosome 4, and will be filtered out of our final cohort because of the noisy coverage in both tumor and normal throughout the region. (b) Multiple loose ends in TCGA glioblastoma multiforme sample 5415. These loose ends will all be filtered out of our final cohort because of the noise, or “waviness”, in the coverage profile. (c) Two loose ends in genome graph for TCGA glioblastoma multiforme sample 0125. These loose ends will be filtered out of our final cohort because the coverage in the paired normal is not flat. (d) A pair of loose ends in TCGA lung squamous cell cancer sample 2756. The coverage ratio shows an apparent gain, due to loss of heterozygosity of a germline copy number loss. These loose ends will be filtered out of our final cohort because of the flat tumor coverage and normal coverage change. (e) Filtering pipeline to evaluate fitted loose ends. *δ*, magnitude change in median coverage value from fused side to unfused side. *β*, sample-specific value related to purity and ploidy indicating magnitude of coverage change corresponding to a single copy state change, as calculated in Eq. S9. *std*, standard deviation (here, the sum of standard deviations of coverage ratio values on the fused and unfused sides of the loose end). *w*, autocorrelation value. Threshold empirically chosen. *ε*, convergence of copy number fit of local subgraph. High values of *ε* indicate an optimal solution of copy number assignments (and thereby loose end incorporation) may not have been reached. *f dr*, false discovery rate using Bonferroni correction. (f) Scatter plot showing estimates of filtering pipeline performance based on blinded visual inspection by five co-authors. TPR, true positive rate. FPR, false positive rate.

**Extended data Fig. 2.**
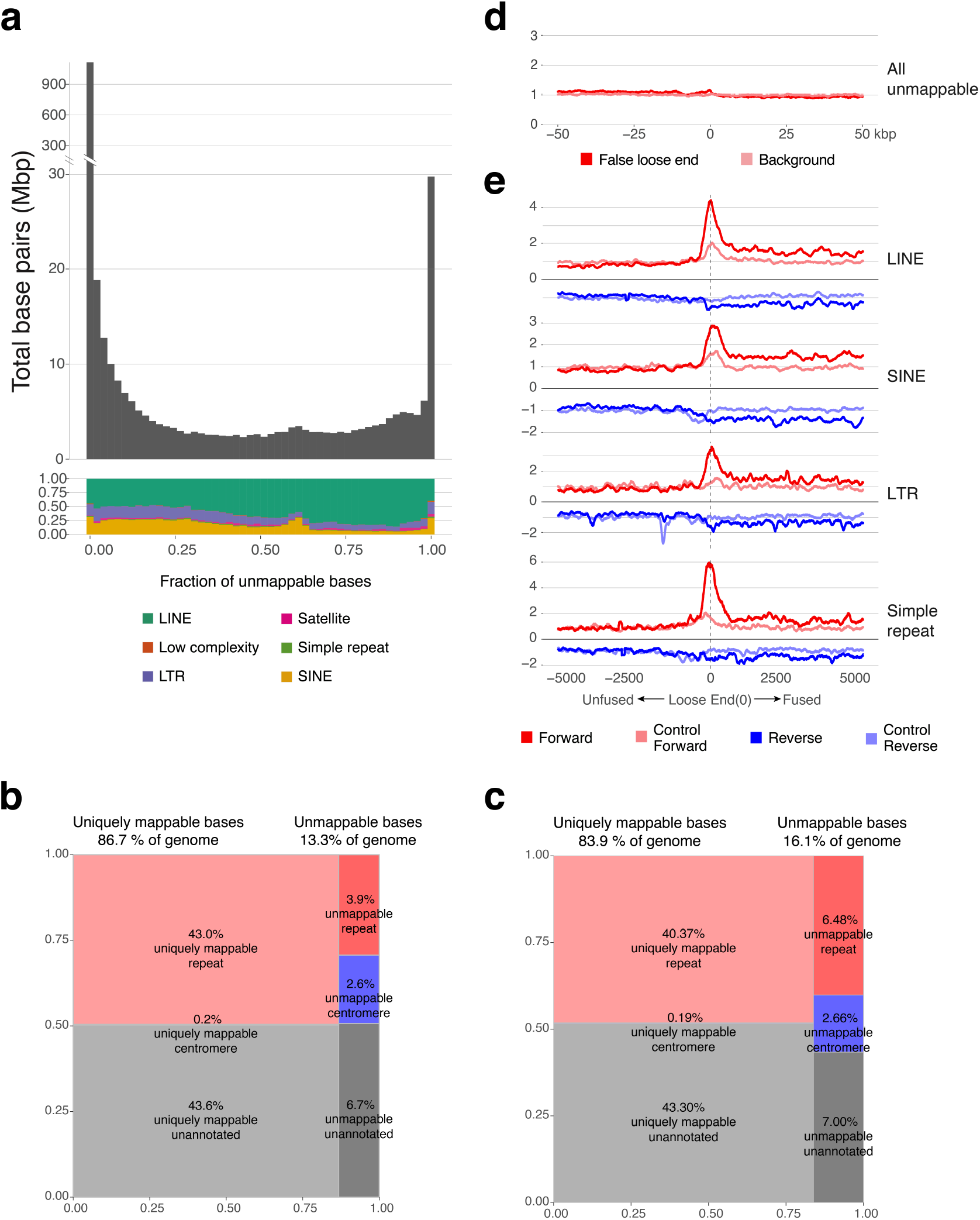
Loose ends are mated to repetitive sequence. (a) Histogram of bases covered by RepeatMasker annotations categorized by the fraction of bases within the annotation with MAPQ < 60 based on simulation with 150bp reads. Above, count of bases falling in any annotated region. Below, breakdown of types of annotation represented in each bin. (b) Composition of hg19 reference genome, based on mappability by 150 bp reads. (c) Composition of hg19 reference genome, based on mappability by 101 bp reads. (d) Aggregate count of all unmappable coordinates surrounding false positive loose ends, normalized to background. Red, aggregate counts around false positive loose ends. Pink, aggregate counts around background sampled coordinates. (e) Normalized aggregate count of loose reads aligning relative to loose breakpoint. Y-axis above zero indicates alignment to loose end-supporting strand; below zero indicates alignment to opposite strand. Top, aggregate count of loose reads whose mates align to LINE annotations. Second from top, aggregate count of loose reads whose mates align to SINE annotations. Third from top, aggregate count of loose reads whose mates align to LTR annotations. Bottom, aggregate count of loose reads whose mates align to simple repeat annotations. Red, aggregate count from loose end-supporting (“forward”) strand in tumor sample. Pink, aggregate count from forward strand in normal sample. Dark blue, aggregate count from opposite (“reverse”) strand in tumor sample. Light blue, aggregate count from reverse strand in normal sample.

**Extended data Fig. 3.**
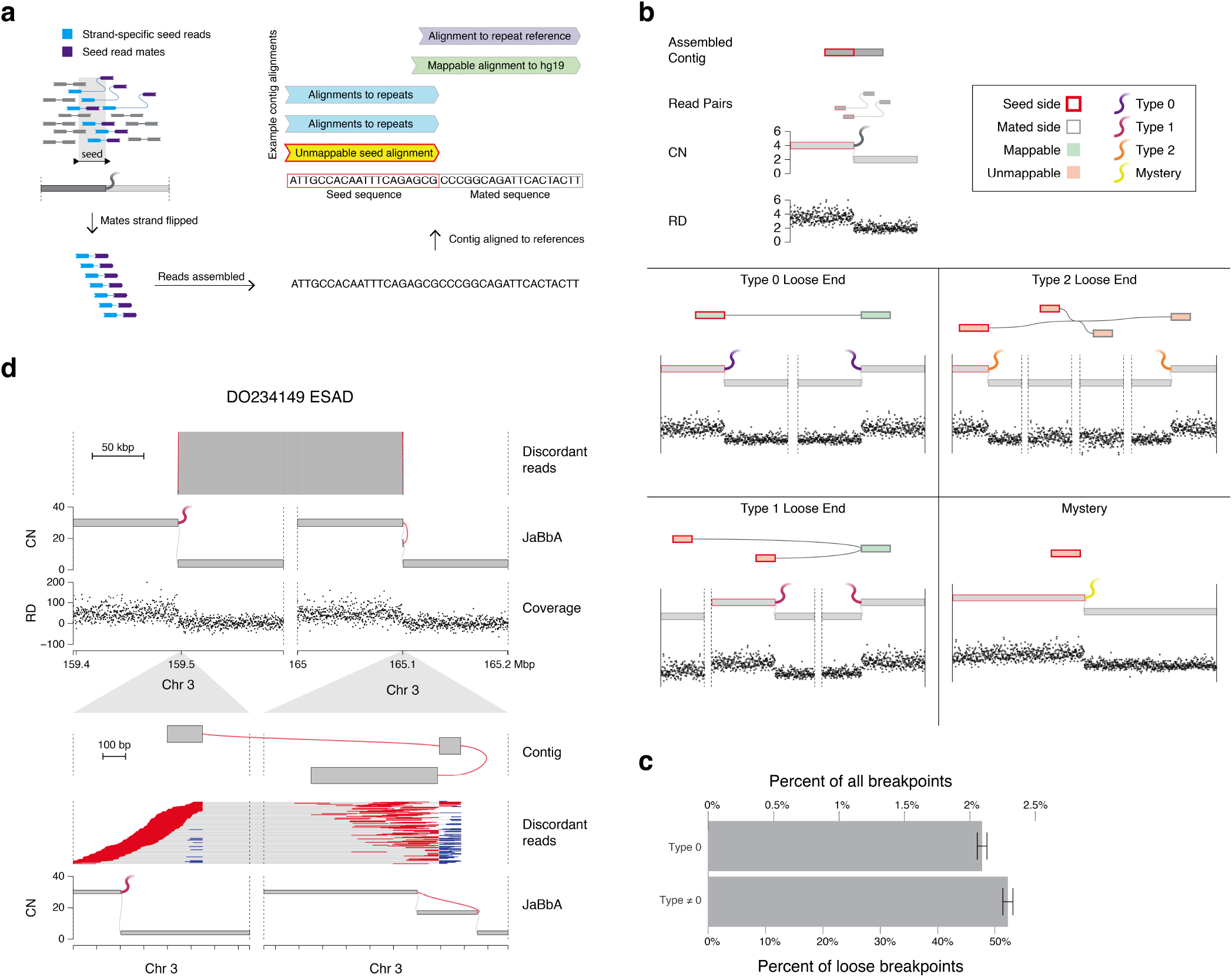
Pipeline to classify loose ends. (a) Schematic of the categorization pipeline. Strand-specific reads aligned to a seed near the loose end and their mates are identified. The read sequences of the seed reads in their original reading frame and the mates reverse complemented from their original reading frame are assembled using Fermi. Assembled contigs are aligned to the human reference, viral references, and databases of repeat sequences. The alignments are parsed to categorize the loose end. (b) Diagram of four categories of underlying loose end explanations. Left: top, diagram of initial loose end. Read pairs align partially as loose reads. Assembling loose read pairs gives a contig constituting the seed (sequence local to the loose end) and the mate (sequence representing the other side of the underlying fusion) Middle, diagram of a Type 0 loose end. The seed and mate sequence align discordantly with MAPQ=60. Bottom, diagram of a Type 1 loose end. The mate sequence may align to a single locus with MAPQ=60, but the seed sequence aligns to multiple loci. Right: middle, diagram of a Type 2 loose end. Both the seed sequence and the mate sequence align to multiple loci. Bottom, diagram of a mystery loose end. No contig is assembled, or the entire contig represents the seed sequence only. (c) Bar plot representing burden of loose ends with respect to all breakpoints (defined as the sum of breakpoints involved in junctions and true loose ends) throughout the cohort, as well as the relative burden of Type 0 loose ends compared to all other types (Type 1, Type 2, Mystery). Error bars show 95% confidence interval. (d) Example Type 1 loose end with complex rearrangement subcategorization from ICGC sample DO234149. A short (< 100 bp) segment of mated sequence is inverted. Some discordant reads do not include alignment to the inverted segment, and some split reads only align to the inverted segment and do not extend through the foldback junction. This complex structure could have caused ambiguity that an SV caller may not be able to resolve. CN, copy number. RD, read depth.

**Extended data Fig. 4.**
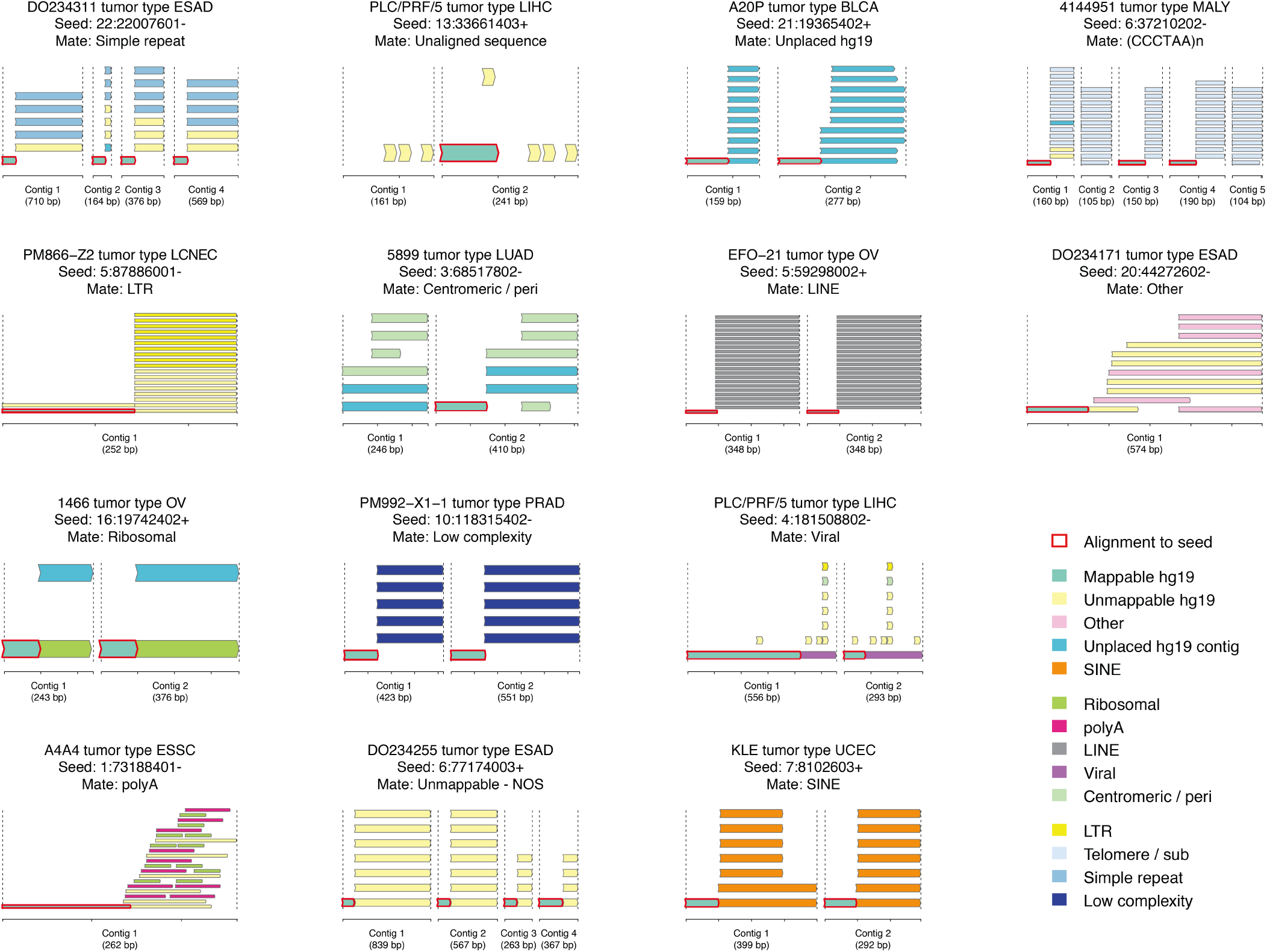
Gallery of mated repeats. For each example, all informative contigs (contigs containing mate sequence) are shown (x-axis is contig coordinates). Each instance of a partial alignment of the contig sequence to a human or repeat reference is indicated with a bar corresponding to the substring of contig sequence aligned. Characteristics of the partial alignments of each contig are indicated by color. The “seed sequence” of every contig is identified as the substring corresponding to an alignment to the seed locus and strand, indicated here with a red outline. Mate sequences with multiple types of repeat alignments follow a heuristic hierarchy to assign the most representative repeat type. All examples shown have mappable seeds and unmappable mates, and therefore belong to the “Type 1 loose end” category.

**Extended data Fig. 5.**
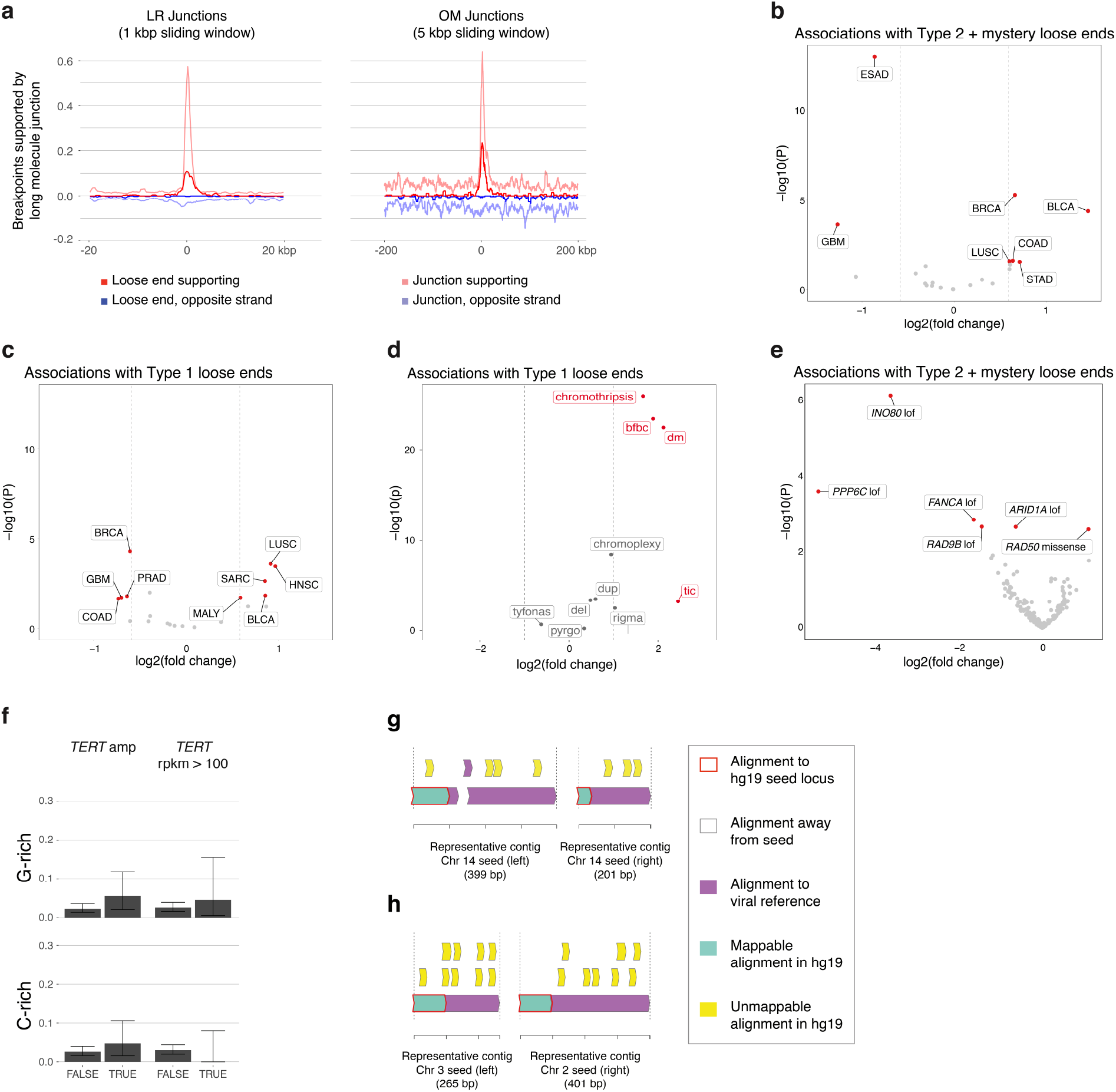
Additional enrichments. (a) Fraction of breakpoints with long range junction call breakpoints in the vicinity. Left, linked read junction calls. Right, optical mapping junction calls. Red, long range junction breakpoints on the same strand as a short read loose end. Dark blue, long range junction breakpoints on the opposite strand as a short read loose end. Pink, long range junction breakpoints on the same strand as a short read junction breakpoint. Light blue, long range junction breakpoints on the opposite strand as a short read junction breakpoint. (b) Volcano plot of associations of Type 2 loose ends with tumor types, based on a generalized linear model. Tumor types with high magnitude of change (*|log*2(*f oldchange*)*| > log*2(1.5)) and high significance (*f dr <* 0.1) are highlighted in red. (c) Volcano plot of associations of Type 1 loose ends with tumor types, based on a generalized linear model. Tumor types with high magnitude of change (*|log*2(*f oldchange*)*| > log*2(1.5)) and high significance (*f dr <* 0.1) are highlighted in red. (d) Volcano plot of associations of Type 1 loose ends with complex SV patters, based on a generalized linear model. Patterns with high magnitude of change (*|log*2(*f oldchange*)*| >* 1) and high significance (*f dr <* 0.1) are highlighted in red. (e) Volcano plot of associations of Type 2 loose ends with genotypes, based on a generalized linear model. Genotypes with high magnitude of change (*|log*2(*f oldchange*)*| > log*2(1.5)) and high significance (*f dr <* 0.1) are highlighted in red. (f) Bar plot of fraction of samples with a given genotype containing at least one telomeric loose end. Error bars show 95% confidence interval. (g-h) Contig alignments of viral loose ends in **Fig. 6d-e**.

**Extended data Fig. 6.**
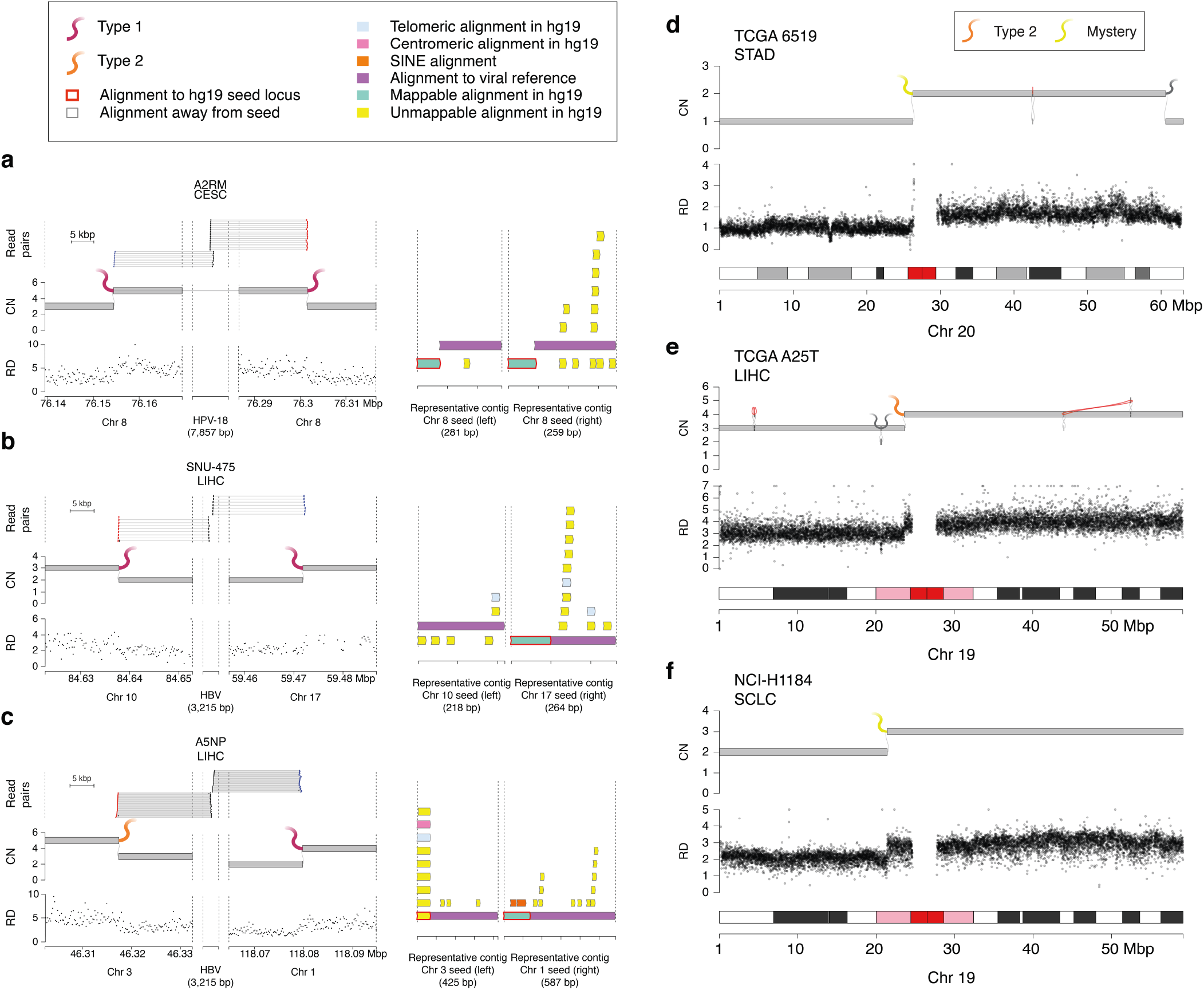
Viral and centromeric loose ends. (a-c) Examples of apparent viral-mediated rearrangements. Left: From top, alignments of read pairs with discordant alignments between human and viral genome. JaBbA graph representation of the sample showing two loose ends corresponding to the discordant read alignments. Normalized binned read depth data, showing corresponding coverage depth changes. Right: Corresponding contig alignments (x-axis is coordinate within contig sequence). (a) Example from TCGA cervical carcinoma sample A2RM. (b) Example from liver hepatocellular carcinoma cell line SNU-475. (c) Example from TCGA liver hepatocellular carcinoma sample A5NP. (d-f) Examples of arm-level loss with an explanatory loose end. Top: JaBbA graph with loose end at centromere. Middle: normalized binned read depth. Bins with outlier coverage values (bottom or top 5 percentile) have been removed. Bottom: cytobands. (d) Loose end within centromere of chromosome 20 in TCGA stomach adenocarcinoma sample 6519, resulting in loss of 20p. This loose end is characterized as a mystery with unmappable centromeric seed sequence. (e) Loose end in peri-centromeric region of chromosome 19 in TCGA liver cancer sample A25T, resulting in loss of 19p. This loose end is characterized as a Type 2 with unmappable peri-centromeric seed sequence mated to unmappable-NOS sequence. (f) Loose end in peri-centromeric region of chromosome 19 in small cell lung cancer cell line NCI-H1184, resulting in loss of 19p. This loose end is characterized as a mystery with unmappable peri-centromeric seed sequence. CN, copy number. RD, read depth.

## References

1. Korbel, J. O. et al. Paired-end mapping reveals extensive structural variation in the human genome. Science 318, 420–426 (2007).

2. Kidd, J. M. et al. Mapping and sequencing of structural variation from eight human genomes. Nature 453, 56–64 (2008).

3. Iakovishina, D., Janoueix-Lerosey, I., Barillot, E., Regnier, M. & Boeva, V. Sv-bay: structural variant detection in cancer genomes using a bayesian approach with correction for gc-content and read mappability. Bioinformatics 32, 984–992 (2016).

4. Yang, R. et al. Integrated analysis of whole-genome paired- end and mate-pair sequencing data for identifying genomic structural variations in multiple myeloma. Cancer informatics 13, CIN–S13783 (2014).

5. Yang, L. et al. Diverse mechanisms of somatic structural variations in human cancer genomes. Cell 153, 919–929 (2013).

6. Sun, R. et al. Breakpointer: using local mapping artifacts to support sequence breakpoint discovery from single-end reads. Bioinformatics 28, 1024–1025 (2012).

7. Marschall, T. et al. Clever: clique-enumerating variant finder. Bioinformatics 28, 2875–2882 (2012).

8. Hayes, M., Pyon, Y. S. & Li, J. A model-based clustering method for genomic structural variant prediction and geno-typing using paired-end sequencing data. PloS one 7, e52881 (2012).

9. Chiara, M., Pesole, G. & Horner, D. S. Svm 2: an improved paired-end-based tool for the detection of small genomic structural variations using high-throughput single-genome resequencing data. Nucleic acids research 40, e145–e145 (2012).

10. Zeitouni, B. et al. Svdetect: a tool to identify genomic structural variations from paired-end and mate-pair sequencing data. Bioinformatics 26, 1895–1896 (2010).

11. Hormozdiari, F., Hajirasouliha, I., McPherson, A., Eichler, E. E. & Sahinalp, S. C. Simultaneous structural variation discovery among multiple paired-end sequenced genomes. Genome research 21, 2203–2212 (2011).

12. Quinlan, A. R. et al. Genome-wide mapping and assembly of structural variant breakpoints in the mouse genome. Genome research 20, 623–635 (2010).

13. Chen, K. et al. Breakdancer: an algorithm for high-resolution mapping of genomic structural variation. Nature methods 6, 677–681 (2009).

14. Sindi, S. S., Önal, S., Peng, L. C., Wu, H.-T. & Raphael, B. J. An integrative probabilistic model for identification of structural variation in sequencing data. Genome biology 13, 1–25 (2012).

15. Ye, K., Schulz, M. H., Long, Q., Apweiler, R. & Ning, Z. Pindel: a pattern growth approach to detect break points of large deletions and medium sized insertions from paired-end short reads. Bioinformatics 25, 2865–2871 (2009).

16. Chen, X. et al. Manta: rapid detection of structural variants and indels for germline and cancer sequencing applications. Bioinformatics 32, 1220–1222 (2016).

17. Bickhart, D. M. et al. Raptr-sv: a hybrid method for the detection of structural variants. Bioinformatics 31, 2084–2090 (2015).

18. Layer, R. M., Chiang, C., Quinlan, A. R. & Hall, I. M. Lumpy: a probabilistic framework for structural variant discovery. Genome biology 15, 1–19 (2014).

19. Hart, S. N. et al. Softsearch: integration of multiple sequence features to identify breakpoints of structural variations. PloS one 8, e83356 (2013).

20. Hayes, M. & Li, J. Bellerophon: a hybrid method for detecting interchromo-somal rearrangements at base pair resolution using next-generation sequencing data 14, 1–9 (2013).

21. Jiang, Y., Wang, Y. & Brudno, M. Prism: pair-read informed split-read mapping for base-pair level detection of insertion, deletion and structural variants. Bioinformatics 28, 2576–2583 (2012).

22. Zhang, J., Wang, J. & Wu, Y. An improved approach for accurate and efficient calling of structural variations with low-coverage sequence data. In BMC bioinformatics, vol. 13, 1–11 (Springer, 2012).

23. Bartenhagen, C. & Dugas, M. Robust and exact structural variation detection with paired-end and soft-clipped alignments: Softsv compared with eight algorithms. Briefings in bioinformatics 17, 51–62 (2016).

24. Wang, J. et al. Crest maps somatic structural variation in cancer genomes with base-pair resolution. Nature methods 8, 652–654 (2011).

25. Suzuki, S., Yasuda, T., Shiraishi, Y., Miyano, S. & Nagasaki, M. Clipcrop: a tool for detecting structural variations with single-base resolution using soft-clipping information 12, 1–9 (2011).

26. Zhang, Z. D. et al. Identification of genomic indels and structural variations using split reads. BMC genomics 12, 1–12 (2011).

27. Grimm, D., Hagmann, J., Koenig, D., Weigel, D. & Borgwardt, K. Accurate indel prediction using paired-end short reads. BMC genomics 14, 1–10 (2013).

28. Barrick, J. E. et al. Identifying structural variation in haploid microbial genomes from short-read resequencing data using breseq. BMC genomics 15, 1–17 (2014).

29. Schröder, J. et al. Socrates: identification of genomic rearrangements in tumour genomes by re-aligning soft clipped reads. Bioinformatics 30, 1064–1072 (2014).

30. Zhang, Z. et al. Sprites: detection of deletions from sequencing data by re-aligning split reads. Bioinformatics 32, 1788–1796 (2016).

31. Wala, J. A. et al. Svaba: genome-wide detection of structural variants and indels by local assembly. Genome research 28, 581–591 (2018).

32. Cameron, D. L. et al. Gridss: sensitive and specific genomic rearrangement detection using positional de bruijn graph assembly. Genome research 27, 2050–2060 (2017).

33. Cameron, D. L. et al. Gridss2: comprehensive characterisation of somatic structural variation using single breakend variants and structural variant phasing (2021). Published online February 16, 2021.

34. Hajirasouliha, I. et al. Detection and characterization of novel sequence insertions using paired-end next-generation sequencing. Bioinformatics 26, 1277–1283 (2010).

35. Iqbal, Z., Caccamo, M., Turner, I., Flicek, P. & McVean, G. De novo assembly and genotyping of variants using colored de bruijn graphs. Nature genetics 44, 226–232 (2012).

36. Rausch, T. et al. Delly: structural variant discovery by integrated paired-end and split-read analysis. Bioinformatics 28, i333–i339 (2012).

37. Abo, R. P. et al. Breakmer: detection of structural variation in targeted massively parallel sequencing data using kmers. Nucleic acids research 43, e19–e19 (2015).

38. Li, Y. et al. Patterns of somatic structural variation in human cancer genomes. Nature 578, 112–121 (2020).

39. Hadi, K. et al. Distinct classes of complex structural variation uncovered across thousands of cancer genome graphs. Cell 183, 197–210 (2020).

40. Davoli, T., Denchi, E. L. & de Lange, T. Persistent telomere damage induces bypass of mitosis and tetraploidy. Cell 141, 81–93 (2010).

41. Thompson, S. L., Bakhoum, S. F. & Compton, D. A. Mechanisms of chromosomal instability. Current biology 20, R285–R295 (2010).

42. Birkbak, N. J. et al. Paradoxical relationship between chromosomal instability and survival outcome in cancer. Cancer research 71, 3447–3452 (2011).

43. Stephens, P. J. et al. Massive genomic rearrangement acquired in a single catastrophic event during cancer development. cell 144, 27–40 (2011).

44. Bakhoum, S. F., Compton, D. A. et al. Chromosomal instability and cancer: a complex relationship with therapeutic potential. The Journal of clinical investigation 122, 1138–1143 (2012).

45. Crasta, K. et al. Dna breaks and chromosome pulverization from errors in mitosis. Nature 482, 53–58 (2012).

46. McGranahan, N., Burrell, R. A., Endesfelder, D., Novelli, M. R. & Swanton, C. Cancer chromosomal instability: therapeutic and diagnostic challenges: ‘exploring aneuploidy: the significance of chromosomal imbalance’review series. EMBO reports 13, 528–538 (2012).

47. Yates, L. R. & Campbell, P. J. Evolution of the cancer genome. Nature Reviews Genetics 13, 795–806 (2012).

48. Orr, B. & Compton, D. A. A double-edged sword: how oncogenes and tumor suppressor genes can contribute to chromosomal instability. Frontiers in oncology 3, 164 (2013).

49. Sansregret, L., Vanhaesebroeck, B. & Swanton, C. Determinants and clinical implications of chromosomal instability in cancer. Nature Reviews Clinical Oncology 15, 139 (2018).

50. Eitan, R. & Shamir, R. Reconstructing cancer karyotypes from short read data: the half empty and half full glass. BMC bioinformatics 18, 1–14 (2017).

51. Ritz, A. et al. Characterization of structural variants with single molecule and hybrid sequencing approaches. Bioinformatics 30, 3458–3466 (2014).

52. Sedlazeck, F. J. et al. Accurate detection of complex structural variations using single-molecule sequencing. Nature methods 15, 461–468 (2018).

53. Huddleston, J. et al. Discovery and genotyping of structural variation from long-read haploid genome sequence data. Genome research 27, 677–685 (2017).

54. Sudmant, P. H. et al. An integrated map of structural variation in 2,504 human genomes. Nature 526, 75–81 (2015).

55. de Koning, A. J., Gu, W., Castoe, T. A., Batzer, M. A. & Pollock, D. D. Repetitive elements may comprise over two-thirds of the human genome. PLoS Genet 7, e1002384 (2011).

56. Tubio, J. M. et al. Extensive transduction of nonrepetitive dna mediated by l1 retrotransposition in cancer genomes. Science 345(2014).

57. Rodriguez-Martin, B. et al. Pan-cancer analysis of whole genomes identifies driver rearrangements promoted by line-1 retrotransposition. Nature genetics 52, 306–319 (2020).

58. Hastings, P. J., Lupski, J. R., Rosenberg, S. M. & Ira, G. Mechanisms of change in gene copy number. Nature Reviews Genetics 10, 551–564 (2009).

59. Chen, J.-M., Cooper, D. N., Férec, C., Kehrer-Sawatzki, H. & Patrinos, G. P. Genomic rearrangements in inherited disease and cancer. In Seminars in cancer biology, vol. 20, 222–233 (Elsevier, 2010).

60. Sishc, B. J. & Davis, A. J. The role of the core non-homologous end joining factors in carcinogenesis and cancer. Cancers 9, 81 (2017).

61. Startek, M. et al. Genome-wide analyses of line–line-mediated nonallelic homologous recombination. Nucleic acids research 43, 2188–2198 (2015).

62. Robberecht, C., Voet, T., Esteki, M. Z., Nowakowska, B. A. & Vermeesch, J. R. Nonallelic homologous recombination between retrotransposable elements is a driver of de novo unbalanced translocations. Genome research 23, 411–418 (2013).

63. Nattestad, M. et al. Complex rearrangements and oncogene amplifications revealed by long-read dna and rna sequencing of a breast cancer cell line. Genome research 28, 1126–1135 (2018).

64. Ardui, S., Ameur, A., Vermeesch, J. R. & Hestand, M. S. Single molecule real-time (smrt) sequencing comes of age: applications and utilities for medical diagnostics. Nucleic acids research 46, 2159–2168 (2018).

65. Spies, N. et al. Genome-wide reconstruction of complex structural variants using read clouds. Nature methods 14, 915–920 (2017).

66. Zheng, G. X. et al. Haplotyping germline and cancer genomes with high-throughput linked-read sequencing. Nature biotechnology 34, 303–311 (2016).

67. Chaisson, M. J. et al. Resolving the complexity of the human genome using single-molecule sequencing. Nature 517, 608–611 (2015).

68. Sakamoto, Y., Sereewattanawoot, S. & Suzuki, A. A new era of long-read sequencing for cancer genomics. Journal of human genetics 65, 3–10 (2020).

69. Pendleton, M. et al. Assembly and diploid architecture of an individual human genome via single-molecule technologies. Nature methods 12, 780–786 (2015).

70. Seo, J.-S. et al. De novo assembly and phasing of a korean human genome. Nature 538, 243–247 (2016).

71. Aganezov, S. et al. Comprehensive analysis of structural variants in breast cancer genomes using single-molecule sequencing. Genome Research 30, 1258–1273 (2020).

72. English, A. C., Salerno, W. J. & Reid, J. G. Pbhoney: identifying genomic variants via long-read discordance and interrupted mapping. BMC bioinformatics 15, 1–7 (2014).

73. Wagner, J. et al. Benchmarking challenging small variants with linked and long reads (2020). Published online July 25, 2020.

74. Imieliński, T. & Lipski, W. Incomplete Information in Relational Databases. Journal of the ACM (JACM) 31, 761–791 (1984).

75. Medvedev, P., Fiume, M., Dzamba, M., Smith, T. & Brudno, M. Detecting copy number variation with mated short reads. Genome research 20, 1613–1622 (2010).

76. Greenman, C. D. et al. Estimation of rearrangement phylogeny for cancer genomes. Genome research 22, 346–361 (2012).

77. Oesper, L., Ritz, A., Aerni, S. J., Drebin, R. & Raphael, B. J. Reconstructing cancer genomes from paired-end sequencing data 13, 1–13 (2012).

78. Li, Y., Zhou, S., Schwartz, D. C. & Ma, J. Allele-specific quantification of structural variations in cancer genomes. Cell systems 3, 21–34 (2016).

79. Dzamba, M. et al. Identification of complex genomic rearrangements in cancers using cougar. Genome research 27, 107–117 (2017).

80. McPherson, A. W. et al. Remixt: clone-specific genomic structure estimation in cancer. Genome biology 18, 1–14 (2017).

81. Olshen, A. B., Venkatraman, E., Lucito, R. & Wigler, M. Circular binary segmentation for the analysis of array-based dna copy number data. Biostatistics 5, 557–572 (2004).

82. Benjamini, Y. & Speed, T. P. Summarizing and correcting the gc content bias in high-throughput sequencing. Nucleic acids research 40, e72–e72 (2012).

83. Deshpande, A., Walradt, T., Hu, Y., Koren, A. & Imielinski, M. Robust foreground detection in somatic copy number data (2019). Published online November 20, 2019.

84. Cameron, D. L., Di Stefano, L. & Papenfuss, A. T. Comprehensive evaluation and characterisation of short read general-purpose structural variant calling software. Nature communications 10, 1–11 (2019).

85. Kosugi, S. et al. Comprehensive evaluation of structural variation detection algorithms for whole genome sequencing. Genome biology 20, 117 (2019).

86. Simpson, J. T. & Durbin, R. Efficient de novo assembly of large genomes using compressed data structures. Genome research 22, 549–556 (2012).

87. Li, H. Exploring single-sample snp and indel calling with whole-genome de novo assembly. Bioinformatics 28, 1838–1844 (2012).

88. Bishara, A. et al. Read clouds uncover variation in complex regions of the human genome. Genome research 25, 1570–1580 (2015).

89. Sedlazeck, F. J., Lee, H., Darby, C. A. & Schatz, M. C. Piercing the dark matter: bioinformatics of long-range sequencing and mapping. Nat. Rev. Genet. 19, 329–346 (2018).

90. Fang, L. et al. Linkedsv for detection of mosaic structural variants from linked-read exome and genome sequencing data. Nature communications 10, 1–15 (2019).

91. Elyanow, R., Wu, H.-T. & Raphael, B. J. Identifying structural variants using linked-read sequencing data. Bioinformatics 34, 353–360 (2018).

92. Maciejowski, J., Li, Y., Bosco, N., Campbell, P. J. & De Lange, T. Chromothripsis and kataegis induced by telomere crisis. Cell 163, 1641–1654 (2015).

93. Pearl, L. H., Schierz, A. C., Ward, S. E., Al-Lazikani, B. & Pearl, F. M. Therapeutic opportunities within the dna damage response. Nature Reviews Cancer 15, 166–180 (2015).

94. Fritsch, O., Benvenuto, G., Bowler, C., Molinier, J. & Hohn, B. The ino80 protein controls homologous recombination in arabidopsis thaliana. Molecular cell 16, 479–485 (2004).

95. Tsukuda, T. et al. Ino80-dependent chromatin remodeling regulates early and late stages of mitotic homologous recombination. DNA repair 8, 360–369 (2009).

96. Neumann, F. R. et al. Targeted ino80 enhances subnuclear chromatin movement and ectopic homologous recombination. Genes & development 26, 369–383 (2012).

97. Challa, K. et al. Damage-induced chromatome dynamics link ubiquitin ligase and proteasome recruitment to histone loss and efficient dna repair. Molecular Cell 81, 811–829 (2021).

98. van Attikum, H., Fritsch, O., Hohn, B. & Gasser, S. M. Recruitment of the ino80 complex by h2a phosphorylation links atp-dependent chromatin remodeling with dna double-strand break repair. Cell 119, 777–788 (2004).

99. Keil, J. M. et al. Symmetric neural progenitor divisions require chromatin-mediated homologous recombination dna repair by ino80. Nature communications 11, 1–15 (2020).

100. Gerhold, C.-B., Hauer, M. H. & Gasser, S. M. Ino80-c and swr-c: guardians of the genome. Journal of molecular biology 427, 637–651 (2015).

101. Benitez, A. et al. Fanca promotes dna double-strand break repair by catalyzing single-strand annealing and strand exchange. Molecular cell 71, 621–628 (2018).

102. Maciejowski, J. & de Lange, T. Telomeres in cancer: tumour suppression and genome instability. Nature reviews Molecular cell biology 18, 175 (2017).

103. Lovejoy, C. A. et al. Loss of atrx, genome instability, and an altered dna damage response are hallmarks of the alternative lengthening of telomeres pathway. PLoS Genet 8, e1002772 (2012).

104. Parfenov, M. et al. Characterization of hpv and host genome interactions in primary head and neck cancers. Proceedings of the National Academy of Sciences 111, 15544–15549 (2014).

105. Park, J. S. et al. Physical status and expression of hpv genes in cervical cancers. Gynecologic oncology 65, 121–129 (1997).

106. Williams, R. Global challenges in liver disease. Hepatology 44, 521–526 (2006).

107. Zapatka, M. et al. The landscape of viral associations in human cancers. Nature genetics 52, 320–330 (2020).

108. Liu, H., Yin, H., Li, G., Li, J. & Wang, X. Aperture: Accurate detection of structural variations and viral integrations in circulating tumor dna using an alignment-free algorithm (2020). Published online December 4, 2020.

109. Zack, T. I. et al. Pan-cancer patterns of somatic copy number alteration. Nature Genetics 45, 1134–1140 (2013).

110. Flint, J. et al. Healing of broken human chromosomes by the addition of telomeric repeats. American journal of human genetics 55, 505 (1994).

111. Wilkie, A. O., Lamb, J., Harris, P. C., Finney, R. D. & Higgs, D. R. A truncated human chromosome 16 associated with *α* thalassaemia is stabilized by addition of telomeric repeat (ttaggg) n. Nature 346, 868–871 (1990).

112. Marzec, P. et al. Nuclear-receptor-mediated telomere insertion leads to genome instability in alt cancers. Cell 160, 913–927 (2015).

113. Sieverling, L. et al. Genomic footprints of activated telomere maintenance mechanisms in cancer. Nature communications 11, 1–13 (2020).

114. Barthel, F. P. et al. Systematic analysis of telomere length and somatic alterations in 31 cancer types. Nature genetics 49, 349–357 (2017).

115. Cesare, A. J. & Reddel, R. R. Alternative lengthening of telomeres: models, mechanisms and implications. Nature reviews genetics 11, 319–330 (2010).

116. de Lange, T. Shelterin-mediated telomere protection. Annual review of genetics 52, 223–247 (2018).

117. Aganezov, S. & Raphael, B. J. Reconstruction of clone-and haplotype-specific cancer genome karyotypes from bulk tumor samples. Genome Research 30, 1274–1290 (2020).

118. Umbreit, N. T. et al. Mechanisms generating cancer genome complexity from a single cell division error. Science 368(2020).

119. Dewhurst, S. M. et al. Structural variant evolution after telomere crisis. Nature communications 12, 1–17 (2021).

120. Collins, R. L. et al. A structural variation reference for medical and population genetics. Nature 581, 444–451 (2020).

121. Barretina, J. et al. The cancer cell line encyclopedia enables predictive modelling of anticancer drug sensitivity. Nature 483, 603–607 (2012).

122. Ghandi, M. et al. Next-generation characterization of the cancer cell line encyclopedia. Nature 569, 503–508 (2019).

123. Li, H. & Durbin, R. Fast and accurate short read alignment with Burrows-Wheeler transform. Bioinformatics 25, 1754–1760 (2009).

124. Nguyen, J. V. Genomic mapping: A statistical and algorithmic analysis of the optical mapping system (University of Southern California, 2010).

125. Anantharaman, T. & Mishra, B. False positives in genomic map assembly and sequence validation. In International Workshop on Algorithms in Bioinformatics, 27–40 (Springer, 2001).

126. Valouev, A., Schwartz, D. C., Zhou, S. & Waterman, M. S. An algorithm for assembly of ordered restriction maps from single dna molecules. Proceedings of the National Academy of Sciences 103, 15770–15775 (2006).

127. Sondka, Z. et al. The COSMIC cancer gene census: describing genetic dysfunction across all human cancers. Nat. Rev. Cancer 18, 696–705 (2018).

